# Survey of the Heritability and Sparse Architecture of Gene Expression Traits Across Human Tissues

**DOI:** 10.1101/043653

**Authors:** Heather E. Wheeler, Kaanan P. Shah, Jonathon Brenner, Tzintzuni Garcia, Keston Aquino-Michaels, GTEx Consortium, Nancy J. Cox, Dan L. Nicolae, Hae Kyung Im

## Abstract

Understanding the genetic architecture of gene expression traits is key to elucidating the underlying mechanisms of complex traits. Here, for the first time, we perform a systematic survey of the heritability and the distribution of effect sizes across all representative tissues in the human body. We find that local h^2^ can be relatively well characterized with 59% of expressed genes showing significant h^2^ (FDR < 0.1) in the DGN whole blood cohort. However, current sample sizes (*n* ≤ 922) do not allow us to compute distal h^2^. Bayesian Sparse Linear Mixed Model (BSLMM) analysis provides strong evidence that the genetic contribution to local expression traits is dominated by a handful of genetic variants rather than by the collective contribution of a large number of variants each of modest size. In other words, the local architecture of gene expression traits is sparse rather than polygenic across all 40 tissues (from DGN and GTEx) examined. This result is confirmed by the sparsity of optimal performing gene expression predictors via elastic net modeling. To further explore the tissue context specificity, we decompose the expression traits into cross-tissue and tissue-specific components using a novel Orthogonal Tissue Decomposition (OTD) approach. Through a series of simulations we show that the cross-tissue and tissue-specific components are identifiable via OTD. Heritability and sparsity estimates of these derived expression phenotypes show similar characteristics to the original traits. Consistent properties relative to prior GTEx multi-tissue analysis results suggest that these traits reflect the expected biology. Finally, we apply this knowledge to develop prediction models of gene expression traits for all tissues. The prediction models, heritability, and prediction performance R^2^ for original and decomposed expression phenotypes are made publicly available (https://github.com/hakyimlab/PrediXcan).

**Author Summary:** Gene regulation is known to contribute to the underlying mechanisms of complex traits. The GTEx project has generated RNA-Seq data on hundreds of individuals across more than 40 tissues providing a comprehensive atlas of gene expression traits. Here, we systematically examined the local versus distant heritability as well as the sparsity versus polygenicity of protein coding gene expression traits in tissues across the entire human body. To determine tissue context specificity, we decomposed the expression levels into cross-tissue and tissue-specific components. Regardless of tissue type, we found that local heritability, but not distal heritability, can be well characterized with current sample sizes. We found that the distribution of effect sizes is more consistent with a sparse local architecture in all tissues. We also show that the cross-tissue and tissue-specific expression phenotypes constructed with our orthogonal tissue decomposition model recapitulate complex Bayesian multi-tissue analysis results. This knowledge was applied to develop prediction models of gene expression traits for all tissues, which we make publicly available.

## Introduction

Regulatory variation plays a key role in the genetics of complex traits [1–3]. Methods that partition the contribution of environment and genetic components are useful tools to understand the biology underlying complex traits. Partitioning heritability into different functional classes (e.g. promoters, coding regions, DNase I hypersensitivity sites) has been successful in quantifying the contribution of different mechanisms that drive the etiology of diseases [3–5].

Most human expression quantitative trait loci (eQTL) studies have focused on how local genetic variation affects gene expression in order to reduce the multiple testing burden that would be required for a global analysis [6, 7]. Furthermore, when both local and distal eQTLs are reported [8–10], effect sizes and replicability are much higher for local eQTLs. While many common diseases are likely polygenic [11–13], it is unclear whether gene expression levels are also polygenic or instead have simpler genetic architectures. It is also unclear how much these expression architectures vary across genes [6].

Bayesian Sparse Linear Mixed Modeling (BSLMM) models complex traits as a mixture of sparse and polygenic contributions. The sparse component consists of a handful of variants of large effect sizes whereas the polygenic component allows for most variants to contribute to the trait albeit with small effect sizes. BSLMM assumes the genotypic effects come from a mixture of two normal distributions and thus is flexible to both polygenic and sparse genetic architectures as well as everything in-between [14]. The model is enforced by sparsity inducing priors on the regression coefficients. BSLMM allows us to directly estimate the sparse and polygenic components of a trait.

As a somewhat independent approach to determine the sparsity and polygenicity of gene expression traits traits, we can look at the relative prediction performance of sparse and polygenic models. For example, if the true genetic architecture of a trait is polygenic, it is natural to expect that polygenic models will predict better (higher predicted vs. observed R^2^) than sparse ones. We assessed the ability of various models, with different underlying assumptions, to predict gene expression in order to understand the underlying genetic architecture of gene expression. For gene expression prediction, we have shown that sparse models such as LASSO (Least Absolute Shrinkage and Selection Operator) perform better than a polygenic score model and that a model that uses the top eQTL variant outperformed the polygenic score but did not do as well as LASSO or elastic net (mixing parameter *α* = 0.5) [15]. These results suggest that for many genes, the genetic architecture is sparse, but not regulated by a single SNP, which is consistent with previous work describing the allelic heterogeneity of gene expression [16–18].

Thus, gene expression traits with sparse architecture should be better predicted with models such as LASSO, which prefers solutions with fewer parameters, each of large effect [19]. Conversely, highly polygenic traits should be better predicted with ridge regression or similarly polygenic models that prefer solutions with many parameters, each of small effect [20–22]. Elastic net [23] is a good multi-purpose model that encompasses both LASSO and ridge regression at its extremes and has been shown to predict well across many complex traits with diverse genetic architectures [24].

Most previous human eQTL studies were performed in whole blood or lymphoblastoid cell lines due to ease of access or culturabilty [8,25,26]. Although studies with a few other tissues have been published, comprehensive coverage of human tissues was not available until the launching of the Genotype-Tissue Expression (GTEx) Project. GTEx aims to examine the genetics of gene expression more comprehensively and has recently published a pilot analysis of eQTL data from 1641 samples across 43 tissues from 175 individuals [27]. Here we use a much larger set of 8555 samples across 53 tissues corresponding to 544 individuals. One of the findings of the pilot analysis was that a large portion of the local regulation of expression traits is shared across multiple tissues. Corroborating this finding, our prediction model built in DGN whole blood showed robust prediction [15] across the nine tissues with the largest sample size from the GTEx Pilot Project [27].

This shared regulation across tissues implies that there is much to be learned from large sample studies of easily accessible tissues. Yet, a portion of gene regulation seems to be tissue dependent [27]. To further investigate the genetic architecture that is common across tissues or specific to each tissue, we use a mixed effects model-based approach termed orthogonal tissue decomposition (OTD) and sought to decouple the cross-tissue and tissue-specific mechanisms in the rich GTEx dataset. We show that the cross-tissue and tissue-specific expression phenotypes constructed with our OTD model reflect the expected biology.

## Results

### Significant local heritability of gene expression in all tissues

We estimated the local and distal heritability of gene expression levels in 40 tissues from the GTEx consortium and whole blood from the Depression Genes and Networks (DGN) cohort. The sample size in GTEx varied from 72 to 361 depending on the tissue, while 922 samples were available in DGN [26]. We used linear mixed models (see Methods) and calculated variances using restricted maximum likelihood (REML) as implemented in GCTA [28]. For the local heritability component, we used common (minor allele frequency > 0.05) variants within 1Mb of the transcription start and end of each protein coding gene, whereas for the distal component, we used common variants outside of the chromosome where the gene was located.

Table 1 summarizes the local heritability estimate results across all tissues. In order to obtain an unbiased estimate of mean h^2^ across genes, we do not constrain the model to only output h^2^ estimates between 0 and 1. Instead, as done previously [10, 29], we allow the h^2^ estimates to be negative when fitting the model and thus refer to it as the unconstrained REML. This approach reduces the standard error of the estimated mean of heritability (law of large numbers). For distal heritability, the errors in the individual heritability estimates were still too large to render a significant mean distal heritability, even in DGN whole blood, the tissue with the largest sample size (S1 Fig). The local component of h^2^ is relatively well estimated in DGN whole blood with 59% of genes (7474 out of 12719) showing FDR < 0.1 (Table 1).

**Table 1.** Estimates of unconstrained local h^2^ across genes within tissues.

| tissue | n | mean $h^2$ (SE) | % FDR<0.1 | num FDR<0.1 | num expressed |
| --- | --- | --- | --- | --- | --- |
| DGN Whole Blood | 922 | 0.149 (0.0039) | 58.8 | 7474 | 12719 |
| Cross-tissue | 450 | 0.062 (0.017) | 20.2 | 2995 | 14861 |
| Adipose - Subcutaneous | 298 | 0.038 (0.028) | 18.8 | 2666 | 14205 |
| Adrenal Gland | 126 | 0.043 (0.018) | 17.5 | 2479 | 14150 |
| Artery - Aorta | 198 | 0.042 (0.024) | 17.2 | 2385 | 13844 |
| Artery - Coronary | 119 | 0.037 (0.048) | 16.5 | 2326 | 14127 |
| Artery - Tibial | 285 | 0.042 (0.02) | 18.4 | 2489 | 13504 |
| Brain - Anterior cingulate cortex (BA24) | 72 | 0.028 (0.036) | 15.4 | 2235 | 14515 |
| Brain - Caudate (basal ganglia) | 100 | 0.037 (0.047) | 17.2 | 2521 | 14632 |
| Brain - Cerebellar Hemisphere | 89 | 0.049 (0.068) | 16.0 | 2294 | 14295 |
| Brain - Cerebellum | 103 | 0.050 (0.06) | 18.3 | 2646 | 14491 |
| Brain - Cortex | 96 | 0.045 (0.057) | 17.7 | 2593 | 14689 |
| Brain - Frontal Cortex (BA9) | 92 | 0.038 (0.077) | 16.6 | 2416 | 14554 |
| Brain - Hippocampus | 81 | 0.037 (0.05) | 16.4 | 2376 | 14513 |
| Brain - Hypothalamus | 81 | 0.017 (0.043) | 15.2 | 2238 | 14759 |
| Brain - Nucleus accumbens (basal ganglia) | 93 | 0.029 (0.04) | 17.1 | 2498 | 14601 |
| Brain - Putamen (basal ganglia) | 82 | 0.032 (0.064) | 17.3 | 2495 | 14404 |
| Breast - Mammary Tissue | 183 | 0.029 (0.037) | 17.0 | 2496 | 14700 |
| Cells - EBV-transformed lymphocytes | 115 | 0.058 (0.1) | 16.6 | 2066 | 12454 |
| Cells - Transformed fibroblasts | 272 | 0.051 (0.031) | 20.0 | 2552 | 12756 |
| Colon - Sigmoid | 124 | 0.033 (0.019) | 16.0 | 2298 | 14321 |
| Colon - Transverse | 170 | 0.036 (0.058) | 16.3 | 2398 | 14676 |
| Esophagus - Gastroesophageal Junction | 127 | 0.032 (0.039) | 16.1 | 2270 | 14125 |
| Esophagus - Mucosa | 242 | 0.042 (0.03) | 17.1 | 2432 | 14239 |
| Esophagus - Muscularis | 219 | 0.039 (0.015) | 17.7 | 2493 | 14047 |
| Heart - Atrial Appendage | 159 | 0.042 (0.03) | 16.8 | 2327 | 13892 |
| Heart - Left Ventricle | 190 | 0.034 (0.034) | 16.5 | 2203 | 13321 |
| Liver | 98 | 0.033 (0.062) | 16.5 | 2230 | 13553 |
| Lung | 279 | 0.032 (0.034) | 17.0 | 2509 | 14775 |
| Muscle - Skeletal | 361 | 0.033 (0.03) | 17.9 | 2301 | 12833 |
| Nerve - Tibial | 256 | 0.052 (0.087) | 20.6 | 2992 | 14510 |
| Ovary | 85 | 0.037 (0.094) | 17.2 | 2418 | 14094 |
| Pancreas | 150 | 0.047 (0.024) | 17.2 | 2398 | 13941 |
| Pituitary | 87 | 0.038 (0.055) | 16.6 | 2527 | 15183 |
| Skin - Not Sun Exposed (Suprapubic) | 196 | 0.041 (0.034) | 17.0 | 2490 | 14642 |
| Skin - Sun Exposed (Lower leg) | 303 | 0.039 (0.016) | 18.4 | 2687 | 14625 |
| Small Intestine - Terminal Ileum | 77 | 0.036 (0.053) | 17.4 | 2591 | 14860 |
| Spleen | 89 | 0.059 (0.061) | 17.0 | 2451 | 14449 |
| Stomach | 171 | 0.032 (0.025) | 16.1 | 2338 | 14531 |
| Testis | 157 | 0.054 (0.044) | 17.8 | 3009 | 16936 |
| Thyroid | 279 | 0.044 (0.066) | 19.4 | 2838 | 14642 |
| Whole Blood | 339 | 0.033 (0.023) | 18.6 | 2260 | 12160 |

Except for DGN Whole Blood, all tissues are from the GTEx Project. Cross-tissue uses derived expression levels from our orthogonal tissue decomposition (OTD) of GTEx data. Mean heritability ( $h^2$ ) is calculated across genes for each tissue. The standard error (SE) is estimated from the $h^2$ estimates obtained when the expression sample labels are shuffled 100 times. The percentage (%) and number (num) of genes with significant $h^2$ estimates (FDR < 0.1) in each tissue are reported. The number of genes in each tissue with mean RPKM > 0.1 (num expressed) is also reported.

It has been shown that local-eQTLs are more likely to be distal-eQTLs of target genes [30]. However, restricting the distal h^2^ estimates to known eQTLs on non-gene chromosomes discovered in a separate cohort (see Methods) did not improve distal h^2^ estimates.

We examined the sensitivity of our local h^2^ estimates to uneven linkage disequilibrium (LD) across the genome using LDAK [31], a method proposed to account for LD. Overall, we find good concordance between GCTA and LDAK estimates, with slightly lower LDAK estimates (S2 Fig). Given the limited sample size we will focus on local regulation for the remainder of the paper.

Across all tissues 15-59% of genes had significant estimates (FDR < 0.1) as shown in Table 1. Gene expression local heritability estimates were consistent between whole blood tissue from the DGN and GTEx cohorts (Spearman's *ρ* = 0.27 [95% confidence interval (CI): 0.25-0.30]). DGN estimates made here were also consistent with those made in independent blood cohorts from previous studies. Comparing DGN to Price et al. [29] and Wright et al. [10], *ρ* = 0.32 [0.30-0.35] and 0.073 [0.047-0.10], respectively. Spearman's *ρ* between Price et al. [29] and Wright et al. [10] gene expression heritability estimates was 0.11 [0.080-0.13].

We also found that more heritable genes tend to be more tolerant to loss of function mutations (Fig. 1). Presumably, such genes are more tolerant to both loss of function mutations and mutations that alter gene expression regulation. This result is consistent with the finding that more eQTLs are found in more tolerant genes [32].

**Figure 1.**
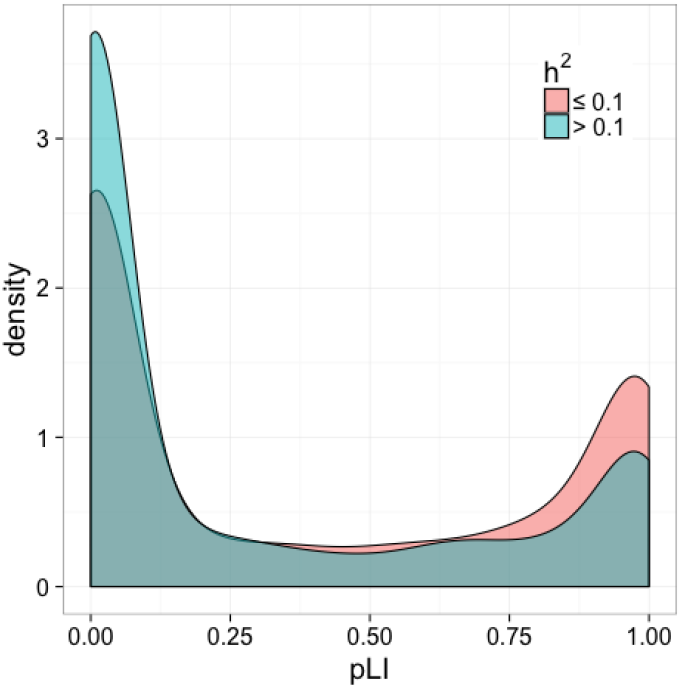
Genes with heritable expression in DGN whole blood are more tolerant to loss of function mutations. The distribution of the probability of being loss-of-function intolerant (pLI) for each gene (from the Exome Aggregation Consortium [32]) dichotomized by local heritability estimates. The Kruskal-Wallis rank sum test revealed a significant difference in the pLI of heritability groups (*χ*^2^ = 234, *P* < 10^−52^). More heritable genes (h^2^ > 0.1 in blue) have lower pLI metrics and are thus more tolerant to mutation than genes with lower h^2^.

### Sparse local architecture revealed by direct estimation using Bayesian Sparse Linear Mixed Modeling (BSLMM)

Next, we sought to determine whether the local common genetic contribution to gene expression is polygenic or sparse. In other words, whether many variants with small effects or a small number of large effects were contributing to expression trait variability. To do this, we used the BSLMM [14] approach, which models the genetic contribution as the sum of a sparse component and a highly polygenic component. The parameter PGE in this model represents the proportion of genetic variance explained by sparse effects. Another parameter, the total variance explained (PVE) by additive genetic variants, is a more flexible Bayesian equivalent of the heritability we have estimated using a linear mixed model (LMM) as implemented in GCTA.

For genes with low heritability, our power to discern between sparse and polygenic architecture is limited. In contrast, for highly heritable genes, the credible sets are tighter and we are able to determine that the sparse component is large. For example, the median PGE was 0.99 [95% CI: 0.94-1] for genes with PVE > 0.50 (Fig. 2A). The median PGE was 0.95 [95% CI: 0.72-0.99] for genes with PVE > 0.1. Fittingly, for most (96.3%) of the genes with heritability > 0.10, the number of SNPs included in the model was below 10.

**Figure 2.**
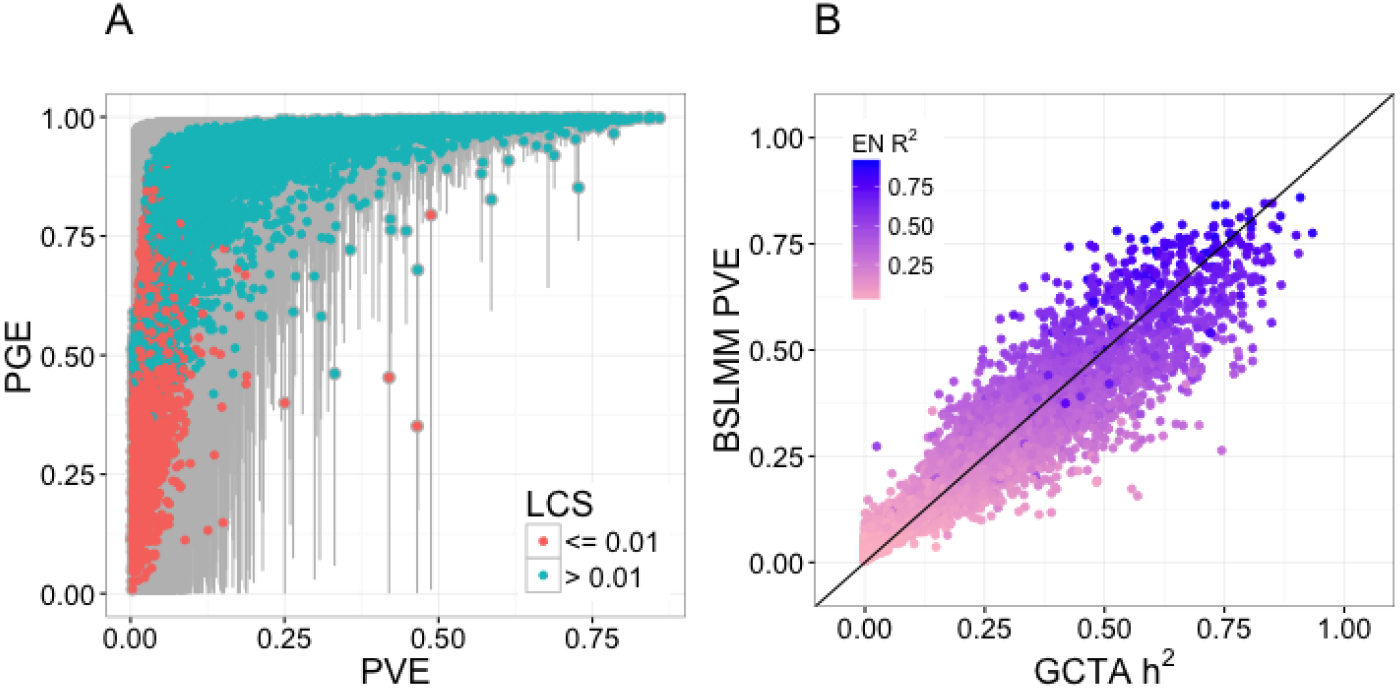
Sparsity estimates using Bayesian Sparse Linear Mixed Models in DGN whole blood. (A) This panel shows a measure of sparsity of the gene expression traits represented by the PGE parameter from the BSLMM approach. PGE is the proportion of the sparse component of the total variance explained by genetic variants, PVE (the BSLMM equivalent of h^2^). The median of the posterior samples of BSLMM output is used as estimates of these parameters. Genes with a lower credible set (LCS) > 0.01 are shown in blue and the rest in red. The 95% credible set of each estimate is shown in gray. For highly heritable genes the sparse component is close to 1, thus for high heritability genes the local architecture is sparse. For lower heritability genes, there is not enough evidence to determine sparsity or polygenicity. (B) This panel shows the heritability estimate from BSLMM (PVE) vs the estimates from GCTA, which are found to be similar (R=0.96). Here, the estimates are constrained to be between 0 and 1 in both models. Each point is colored according to that gene's elastic net *α* = 1 cross-validated prediction correlation squared (EN R^2^). Note genes with high heritability have high prediction R^2^, as expected.

### Prediction models with a sparse component outperform polygenic models

To further confirm the local sparsity of gene expression traits, we looked at the prediction performance of a range of models with different degrees of polygenicity, such as the elastic net model with mixing parameter values ranging from 0 (fully polygenic, ridge regression) to 1 (sparse, LASSO). We performed 10-fold cross-validation using the elastic net [23] to test the predictive performance of local SNPs for gene expression across a range of mixing parameters (*α*). Given common folds and seeds, the mixing parameter that yields the largest cross-validation R^2^ informs the degree of sparsity of each gene expression trait. That is, at one extreme, if the optimal *α* = 0 (equivalent to ridge regression), the gene expression trait is highly polygenic, whereas if the optimal *α* = 1 (equivalent to LASSO), the trait is highly sparse. We found that for most gene expression traits, the cross-validated R^2^ was smaller for *α* = 0 and *α* = 0.05, but nearly identical for *α* = 0.5 through *α* = 1 in the DGN cohort (Fig. 3). An *α* = 0.05 was also clearly suboptimal for gene expression prediction in the GTEx tissues, while models with *α* = 0.5 or 1 had similar predictive power (S3 Fig). Together with the BSLMM results, this suggests that for most genes, the effect of local common genetic variation on gene expression is sparse rather than polygenic.

**Figure 3.**
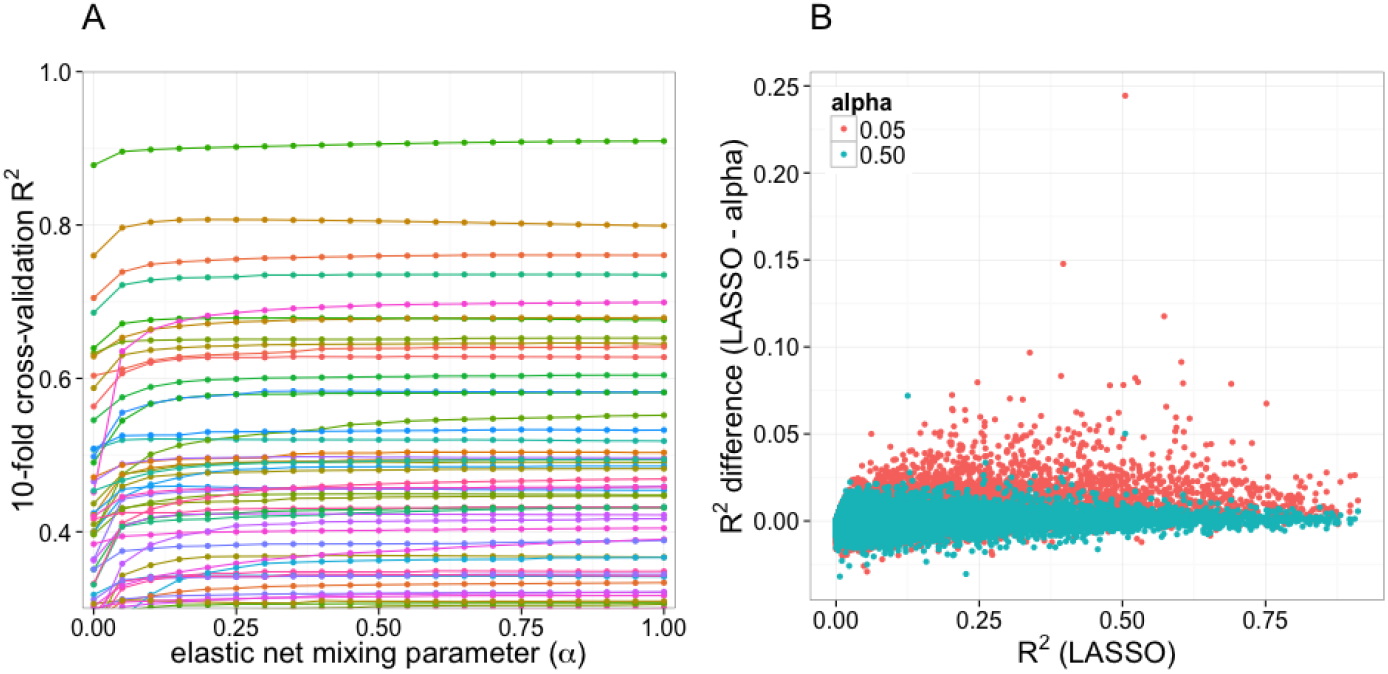
DGN cross-validated predictive performance across the elastic net. Elastic net prediction models were built in the DGN whole blood and performance was quantified by the cross-validated R^2^ between observed and predicted expression levels. (A) This panel shows the 10-fold cross validated R^2^ for 51 genes with R^2^ > 0.3 from chromosome 22 as a function of the elastic net mixing parameters (*α*). Smaller mixing parameters correspond to more polygenic models while larger ones correspond to more sparse models. Each line represents a gene. The performance is in general flat for most values of the mixing parameter except very close to zero where it shows a pronounced dip. Thus polygenic models perform more poorly than sparse models. (B) This panel shows the difference between the cross validated R^2^ of the LASSO model and the elastic net model mixing parameters 0.05 and 0.5 for autosomal protein coding genes. Elastic net with *α* = 0.5 values hover around zero, meaning that it has similar predictive performance to LASSO. The R^2^ difference of the more polygenic model (elastic net with *α* = 0.05) is mostly above the 0 line, indicating that this model performs worse than the LASSO model.

### BSLMM outperforms LMM in estimating h^2^ for small samples

In DGN, there is a strong correlation between BSLMM-estimated PVE and GCTA-estimated h^2^ (Fig. 2B, R=0.96). In contrast, when we applied BSLMM to the GTEx data, we found that many genes had measurably larger BSLMM-estimated PVE than LMM-estimated h^2^ (Fig. 4). This is further confirmation of the predominantly sparse local architecture of gene expression traits: the underlying assumption in the LMM approach to estimate heritability is that the genetic effect sizes are normally distributed, i.e. most variants have small effect sizes. LMM is quite robust to departure from this assumption, but only when the sample size is rather large (S4 Fig). For the relatively small sample sizes in GTEx (*n* ≤ 361), we found that directly modeling the sparse component with BSLMM (polygenic + sparse components) outperforms LMM (single polygenic component) for estimating h^2^. Here, unlike in previous sections (see Table 1 and S1 Fig), both the BSLMM and LMM model estimates are constrained to be between 0 and 1.

**Figure 4.**
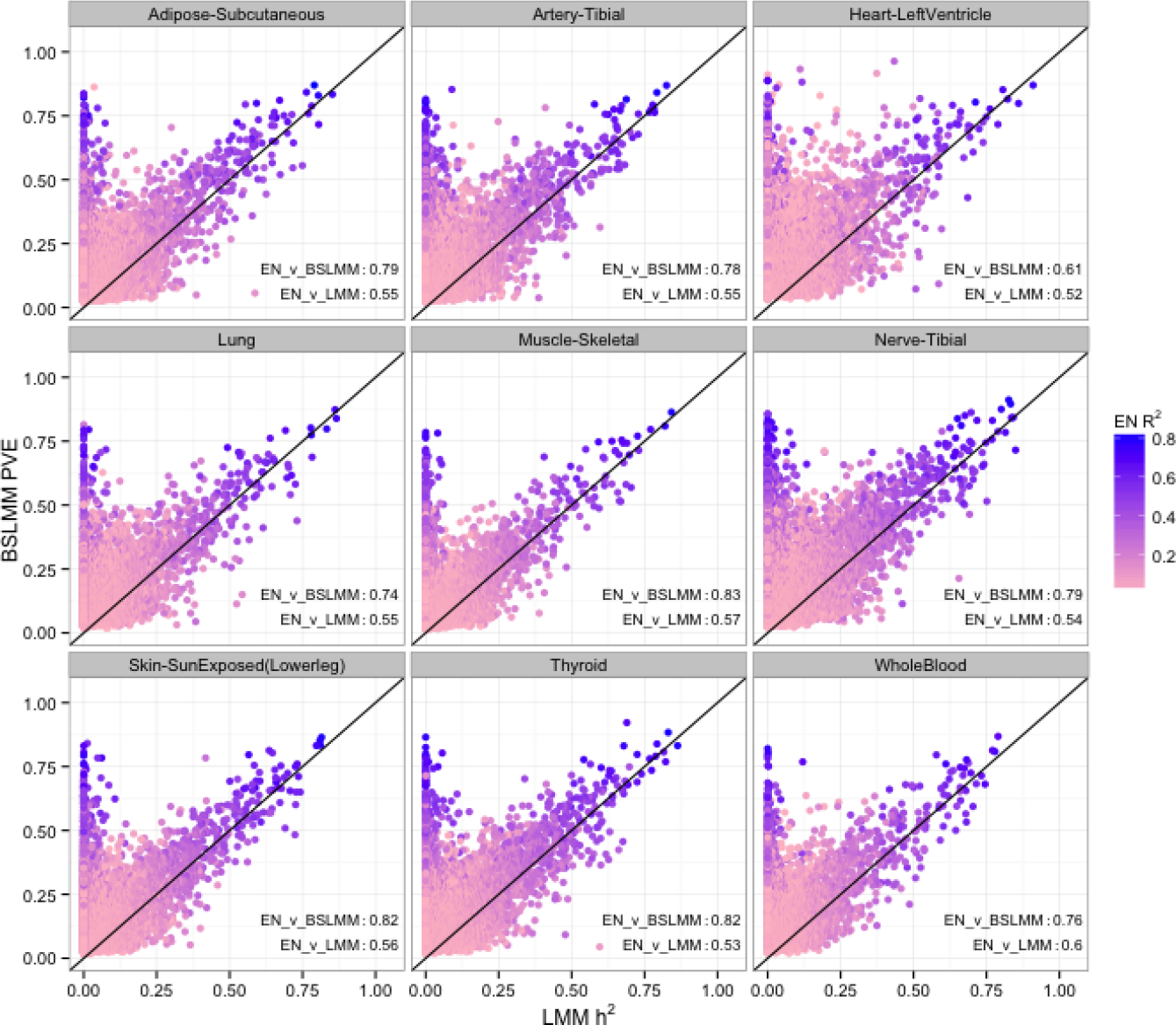
BSLMM vs LMM estimates of heritability in GTEx. This figure shows the comparison between estimates of heritability using BSLMM vs. LMM (GCTA) for GTEx data. Here, in both models the estimates are constrained to be between 0 and 1. For most genes BSLMM estimates are larger than LMM estimates reflecting the fact that BSLMM yields better estimates of heritability because of its ability to account for the sparse component. Each point is colored according to that gene's prediction R2 (correlation squared between cross-validated elastic net prediction vs observed expression denoted EN R^2^). At the bottom left of each panel, we show the correlation between BSLMM (EN_v_BSLMM) and LMM (EN_v_LMM). BSLMM is consistently more correlated with the elastic net correlation. This provides further indication that the local architecture is predominantly sparse.

To ensure that the high values of heritability estimated by BSLMM are due to actually higher h^2^ and not to limitations or bias of the BSLMM, we use the fact that cross-validated prediction R^2^ (observed vs predicted squared correlation when prediction parameters are estimated without including the observations to be predicted) is bounded by the heritable component. In other words, genetically based prediction models can only explain or predict the portion that is heritable. As shown in Figure 4, most of the genes with high BSLMM heritability but low LMM h^2^ have large values of prediction R^2^. We further validated this behavior by showing Price et al. [29] identity-by-descent (IBD) heritability estimates correlate more strongly with BSLMM than LMM estimates (S5 Fig). For all tissues, correlation is higher between IBD estimates and BSLMM than between IBD and LMM. These findings provide strong evidence that LMM is underestimating h^2^ for these genes.

### Orthogonal decomposition of cross-tissue and tissue-specific expression traits

Since a substantial portion of local regulation was shown to be common across multiple tissues [27], we sought to decompose the expression levels into a component that is common across all tissues and tissue-specific components. Figure 5 shows a diagram describing the decomposition for which we use a linear mixed effects model with a person-level random effect (see Methods). We use the posterior mean of this random effect as an estimate of the cross-tissue component. We consider the residual component of this model as the tissue-specific component. We call this approach orthogonal tissue decomposition (OTD) because the cross-tissue and tissue-specific components are assumed to be independent in the model.

**Figure 5.**
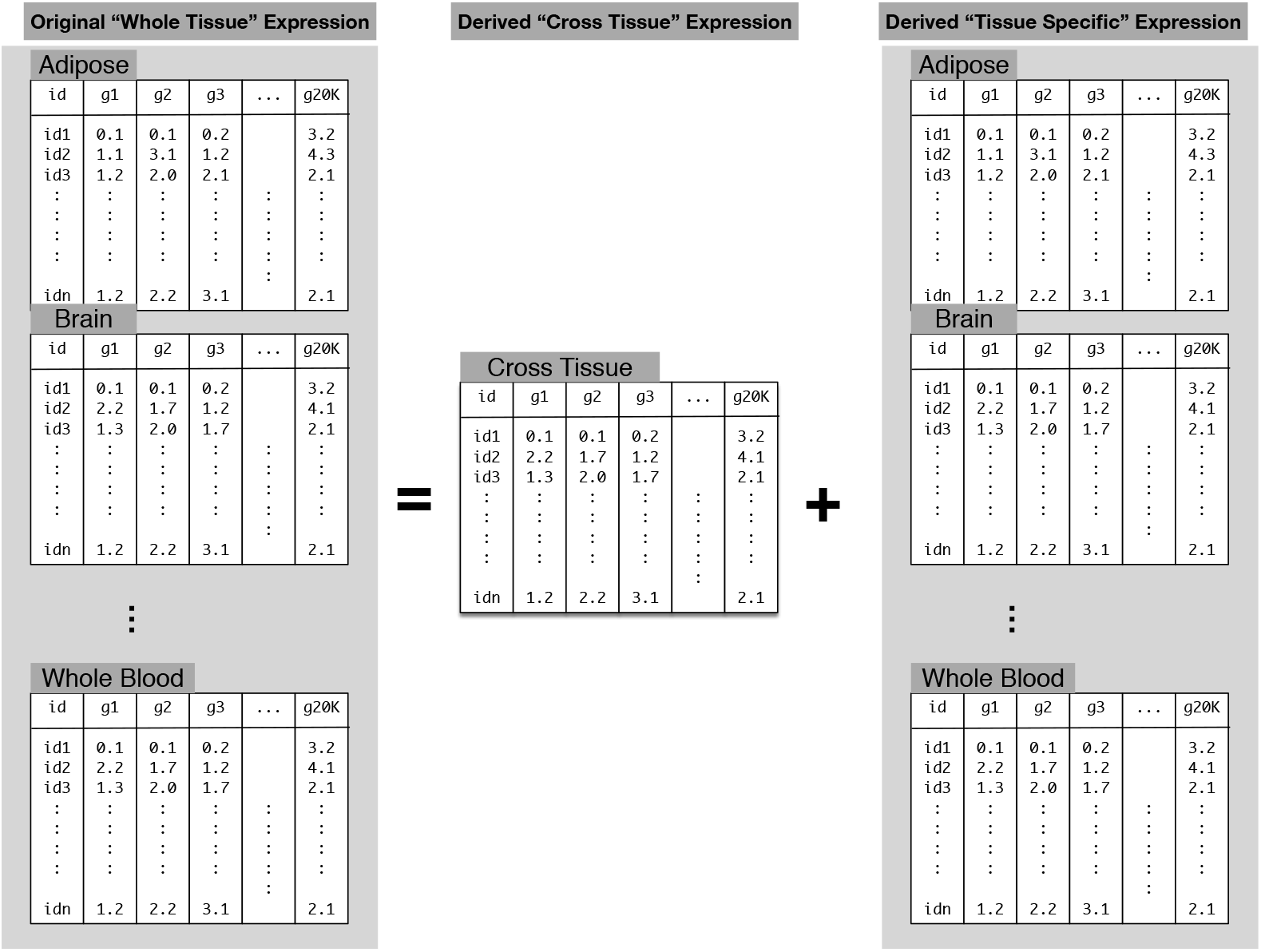
Orthogonal Tissue Decomposition of gene expression traits. For a given gene, the expression level is decoupled into a component that is specific to the individual and another component that is specific to the individual and tissue. The left side of the equation in the figure corresponds to the original “whole tissue” expression levels. The right side has the component specific for the individual, independent of the tissue and the tissue-specific component. Given the lack of multiple replications for a given tissue/individual we use a mixed effects model with a random effect that is specific to the individual. The cross-tissue component is estimated as the posterior mean of the subject-specific random effect. The tissue-specific component is estimated as the residual of the model fit, i.e. the difference between the “whole tissue” expression and the cross-tissue component. The rationale is that once we remove the component that is common across tissues, the remaining will be specific to the tissue. Models are fit one gene at a time. Covariates are not shown to simplify the presentation.

First, we sought to demonstrate that OTD is able to identify the cross-tissue and tissue-specific components via simulations. We generated simulated traits so that the true cross-tissue and tissue-specific components are known. To preserve the correlation structure between genes and tissues, we considered the estimated cross-tissue and tissue-specific components as the true values. Then, we simulated expression traits as the sum of the cross-tissue and tissue-specific components and an error term (expression trait = cross-tissue + tissue-specific + noise). We simulated expression traits using different magnitudes of the noise level and then applied our OTD approach. Simulated values were constructed using noise variance equal in magnitude to the original variance of the expression levels of each gene. This is a rather conservative choice since the variance of the noise component is unlikely to be larger than the total variance of the expression trait. In S6 Fig, we show for one simulated example the correlation between true and OTD-estimated components for one representative gene, *SLTM*, and the distribution of the correlation between true and OTD-estimates for all genes. The median Pearson correlation between true and OTD-estimted cross-tissue components was 0.54 [95% CI: 0.33-0.77] and for tissue-specific components was 0.75 [0.46-0.88], demonstrating the robustness of our model (S6 Fig).

For the OTD derived cross-tissue and 40 tissue-specific expression phenotypes, we computed the local heritability and generated prediction models. The decomposition is applied at the expression trait level so that the downstream genetic regulation analysis is performed separately for each derived trait, cross-tissue and tissue-specific expression, which greatly reduces computational burden.

### Cross-tissue expression phenotype is less noisy and shows higher predictive performance

Our estimates of h^2^ for cross-tissue expression traits are larger than the corresponding estimates for each whole tissue (S7 Fig) because our OTD approach increases the ratio of the genetically regulated component to noise by averaging across multiple tissues. In addition to the increased h^2^, we observe reduction in standard errors of the estimated cross-tissue h^2^. This is due, in part, to the larger effective sample size for cross-tissue phenotypes. There were 450 samples for which cross-tissue traits were available whereas the maximum sample size for whole tissue phenotypes was 362. Similarly, cross-tissue BSLMM PVE estimates had lower error than whole tissue PVE (S8 Fig, S9 Fig).

As for the tissue-specific components, the cross-tissue heritability estimates were also larger and the standard errors were smaller reflecting the fact that a substantial portion of regulation is common across tissues (S10 Fig). The percentage of GCTA h^2^ estimates with FDR <0.1 was much larger for cross-tissue expression (20%) than the tissue-specific expressions (6-13%, S1 Table). Similarly, the percentage of BSLMM PVE estimates with a lower credible set greater than 0.01 was 49% for cross-tissue expression, but ranged from 24-27% for tissue-specific expression (S9 Fig).

Cross-tissue predictive performance exceeded that of both tissue-specific and whole tissue expression as indicated by higher cross-validated R^2^ (S11 Fig). Like whole tissue expression, cross-tissue and tissue-specific expression showed higher predictive performance when using more sparse models. In other words elastic-net models with *α* ≥ 0.5 predicted better than the ones with *α* = 0.05 (S11 Fig).

### Cross-tissue expression phenotype recapitulates published multi-tissue eQTL results

To verify that the cross-tissue phenotype has the properties we expect, we compared our OTD results to those from a joint multi-tissue eQTL analysis [33], which was previously performed on a subset of the GTEx data [27] covering 9 tissues. In particular, we used the posterior probability of a gene being actively regulated (PPA) in a tissue. These analysis results are available on the GTEx portal (see Methods).

First, we reasoned that genes with high cross-tissue h^2^ would be actively regulated in most tissues so that the PPA of a gene would be roughly uniform across tissues. By contrast, a gene with tissue-specific regulation would have concentrated posterior probability in one or a few tissues. Thus we decided to define a measure of uniformity of the posterior probability vector across the 9 tissues using the concept of entropy. More specifically, for each gene we normalized the vector of posterior probabilities so that the sum equaled 1. Then we applied the usual entropy definition (negative of the sum of the log of the posterior probabilities weighted by the same probabilities, see Methods). In other words, we defined a uniformity statistic that combines the nine posterior probabilities into one value such that higher values mean the gene regulation is more uniform across all nine tissues, rather than in just a small subset of the nine.

Thus, we expected that genes with high cross-tissue heritability would show high probability of being active in multiple tissues and have high uniformity measure. Reassuringly, this is exactly what we find. Genes with high cross-tissue heritability concentrate on the higher end of the uniformity measure (Fig. 6, S12 Fig).

**Figure 6.**
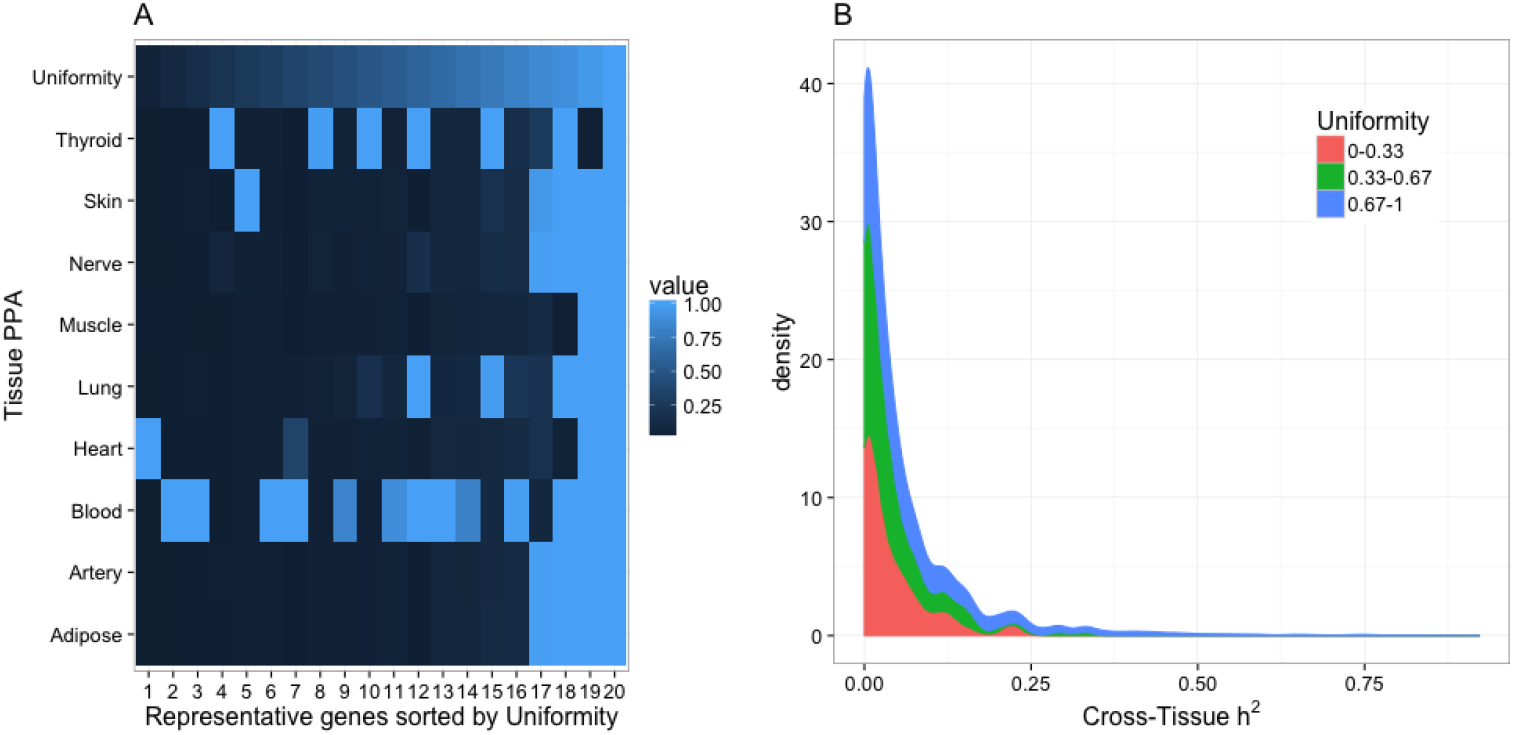
Measure of uniformity of the posterior probability of active regulation vs. cross-tissue heritability. Uniformity was computed using the posterior probability of a gene being actively regulated in a tissue, PPA, from the Flutre et al. [33] multi-tissue eQTL analysis. **(A)** Representative examples showing that genes with PPA concentrated in one tissue were assigned small values of the uniformity measure whereas genes with PPA uniformly distributed across tissues were assigned high value of uniformity measure. See Methods for the entropy-based definition of uniformity. **(B)** This panel shows the distribution of heritability of the cross-tissue component vs. a measure of uniformity of genetic regulation across tissues. The Kruskal-Wallis rank sum test revealed a significant difference in the cross-tissue h^2^ of uniformity groups (*χ*^2^ = 31.4, *P* < 10^−6^).

For the original whole tissue, we expected the whole tissue expression heritability to correlate with the posterior probability of a gene being actively regulated in a tissue. This is confirmed in Figure 7A where PPA in each tissue is correlated with the BSLMM PVE of the expression in that tissue. In the off diagonal elements we observe high correlation between tissues, which was expected given that large portion of the regulation has been shown to be common across tissues. Whole blood has the lowest correlation consistent with whole blood clustering aways from other tissues [27]. In contrast, Figure 7B shows that the tissue-specific expression PVE correlates well with matching tissue PPA but the off diagonal correlations are substantially reduced consistent with these phenotypes representing tissue-specific components. Again whole blood shows a negative correlation which could be indicative of some over correction of the cross-tissue component. Overall these results indicate that the cross-tissue and tissue-specific phenotypes have properties that are consistent with the intended decomposition.

**Figure 7.**
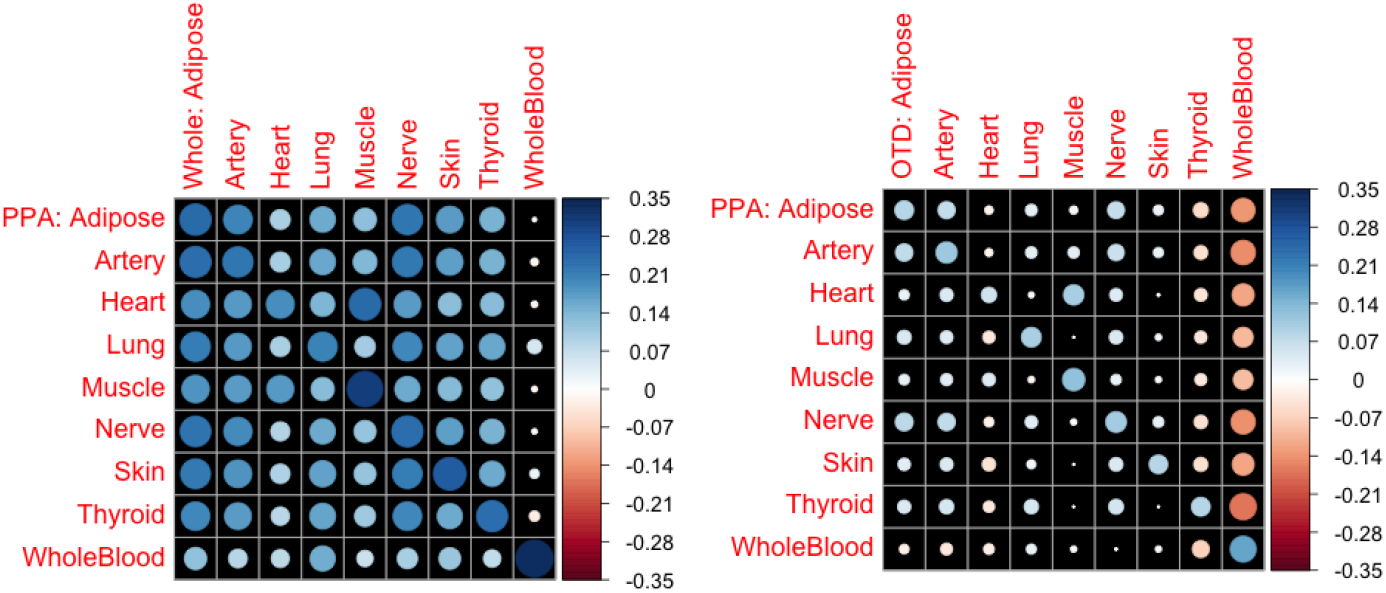
Comparison of heritability of whole tissue or tissue-specific components vs. PPA. Panel (A) of this figure shows the Pearson correlation (R) between the BSLMM PVE of the original (we are calling whole here) tissue expression levels vs. the probability of the tissue being actively regulated in a given tissue (PPA). Matching tissues show, in general, the largest correlation values but most of the off diagonal correlations are also relatively high consistent with the shared regulation across tissues. Panel (B) shows the Pearson correlation between the PVE of the tissue-specific component of expression via orthogonal tissue decomposition (OTD) vs. PPA. Correlations are in general lower but matching tissues show the largest correlation. Off diagonal correlations are reduced substantially consistent with properties that are specific to each tissue. Area of each circle is proportional to the absolute value of R.

## Discussion

Motivated by the key role that regulatory variation plays in the genetic control of complex traits [1–3], we performed a survey of the heritability and patterns of effect sizes of gene expression traits across a comprehensive set of human tissues. We quantified the local and distal heritability of gene expression in DGN and 40 different tissues from the GTEx consortium. Distal components estimates were too noisy to draw meaningful conclusions.

Using results implied by the improved predictive performance of sparse models and by directly estimating sparsity using BSLMM (Bayesian Sparse Linear Mixed Model), we show evidence that local common variant regulation is sparse across all the tissues analyzed here. Our finding that a handful of genetic variants seem to contribute to the variability in gene expression traits has important implications for the strategies used in future investigations of gene regulation and downstream effects on complex traits. For example, most fine mapping methods that attempt to find the causal SNPs that drive a GWAS locus focus on a limited number of variants. This only makes sense in cases where the underlying genetic architecture is sparse, that is, when a handful of causal variants are determining the variability of the traits. We note that any rare variants not tagged by common variants are not included in our gene expression heritability estimates and prediction models. For genes with moderate and low heritability the evidence is not as strong, due to reduced power, but results are consistent with a sparse local architecture. Sparse models capture the most variance in gene expression at current sample sizes and have been used successfully in gene expression prediction methods [15, 36–38]. Better methods to correct for hidden confounders that do not dilute distal signals and larger sample sizes will be needed to determine the properties of distal regulation.

Given that a substantial portion of local regulation is shared across tissues, we proposed here to decompose the expression traits into cross-tissue and tissue-specific components. This approach, called orthogonal tissue decomposition (OTD), aims to decouple the shared regulation from the tissue-specific regulation. We examined the genetic architecture of these derived traits and find that they follow similar patterns to the original whole tissue expression traits. The cross-tissue component benefits from an effectively larger sample size than any individual tissue trait, which is reflected in more accurate heritability estimates and consistently higher prediction performance. Encouragingly, we find that genes with high cross-tissue heritability tend to be regulated more uniformly across tissues. As for the tissue-specific expression traits, we found that they recapitulate correlation with the vector of probability of tissue-specific regulation. Many groups have proposed integrating genotype and expression data to understand complex traits [15,22,37–40]. Through integration of our OTD expression traits with studies of complex diseases, we expect results from the cross-tissue models to relate to mechanisms that are shared across multiple tissues, whereas results from the tissue-specific models will inform us about the context specific mechanisms.

In this paper, we quantitate the genetic architecture of gene expression and develop predictors across tissues. We show that local heritability can be accurately estimated across tissues, but distal heritability cannot be reliably estimated at current sample sizes. Using two different approaches, BSLMM and the elastic net, we show that for common local gene regulation, the genetic architecture is mostly sparse rather than polygenic. Using new expression phenotypes generated in our OTD model, we show that cross-tissue predictive performance exceeded that of both tissue-specific and whole tissue expression as indicated by higher elastic net cross-validated R^2^. Predictors, heritability estimates and cross-validation statistics generated in this study of gene expression architecture are freely available (https://github.com/hakyimlab/PrediXcan) for use in future studies of complex trait genetics.

## Methods

### Genomic and Transcriptomic Data

#### DGN Dataset

We obtained whole blood RNA-seq and genome-wide genotype data for 922 individuals from the Depression Genes and Networks (DGN) cohort [26], all of European ancestry. For all analyses, we used the HCP (hidden covariates with prior) normalized gene-level expression data used for the *trans*-eQTL analysis in Battle et al. [26] and downloaded from the NIMH repository. The 922 individuals were unrelated (all pairwise 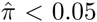) and thus all included in downstream analyses. Imputation of approximately 650K input SNPs (minor allele frequency [MAF] > 0.05, Hardy-Weinberg Equilibrium [P > 0.05], non-ambiguous strand [no A/T or C/G SNPs]) was performed on the Michigan Imputation Server (https://imputationserver.sph.umich.edu/start.html) [41, 42] with the following parameters: 1000G Phase 1 v3 ShapeIt2 (no singletons) reference panel, SHAPEIT phasing, and EUR population. Approximately 1.9M non-ambiguous strand SNPs with MAF > 0.05, imputation R^2^ > 0.8 and, to reduce computational burden, inclusion in HapMap Phase II were retained for subsequent analyses.

#### GTEx Dataset

We obtained RNA-seq gene expression levels from 8555 tissue samples (53 unique tissue types) from 544 unique subjects in the GTEx Project [27] data release on 2014-06-13. Of the individuals with gene expression data, genome-wide genotypes (imputed with 1000 Genomes) were available for 450 individuals. While all 8555 tissue samples were used in the OTD model (described below) to generate cross-tissue and tissue-specific components of gene expression, we used the 40 tissues with the largest sample sizes (*n* > 70) when quantifying tissue-specific effects (see S1 Table). Approximately 2.6M non-ambiguous strand SNPs included in HapMap Phase II were retained for subsequent analyses.

#### Framingham Expression Dataset

We obtained exon array expression and genotype array data from 5257 individuals from the Framingham Heart Study [43]. The final sample size after quality control was 4286. We used the Affymetrix power tools (APT) suite to perform the preprocessing and normalization steps. First the robust multi-array analysis (RMA) protocol was applied which consists of three steps: background correction, quantile normalization, and summarization [44]. The background correction step uses antigenomic probes that do not match known genome sequences to adjust the baseline for detection, and is applied separately to each array. Next, the normalization step utilizes a ‘sketch’ quantile normalization technique instead of a memory-intensive full quantile normalization. The benefit is a much lower memory requirement with little accuracy trade-off for large sample sets such as this one. Next, the adjusted probe values were summarized (by the median polish method) into log-transformed expression values such that one value is derived per exon or gene. The summarized expression values were then annotated more fully using the annotation databases contained in the huex10stprobeset.db (exon-level annotations) and huex10sttranscriptcluster.db (gene-level annotations) R packages available from Bioconductor [45,46]. In both cases gene annotations were provided for each feature.

Plink [47] was used for data wrangling and cleaning steps. The data wrangling steps included updating probe IDs, unifying data to the positive strand, and updating locations to GRCh37. The data cleaning steps included a step to filter for variant and subject missingness and minor alleles, one to filter variants with Hardy-Weinberg exact test, and a step to remove unusual heterozygosity. To make our data compatible with the Haplotype Reference Consortium (HRC) panel (http://www.well.ox.ac.uk/~wrayner/tools/), we used the HRC-check-bin tool in order to carry out data wrangling steps required. The data were then split by chromosome and pre-phased with SHAPEIT [48] using the 1000 Genomes phase 3 panel and converted to vcf format. These files were then submitted to the Michigan Imputation Server (https://imputationserver.sph.umich.edu/start.html) [41, 42] for imputation with the HRC version 1 panel [49]. We applied Matrix eQTL [50] to the normalized expression and imputed genotype data to generate prior eQTLs for our heritability analysis.

### Partitioning local and distal heritability of gene expression

Motivated by the observed differences in regulatory effect sizes of variants located in the vicinity of the genes and distal to the gene, we partitioned the proportion of gene expression variance explained by SNPs in the DGN cohort into two components: local (SNPs within 1Mb of the gene) and distal (eQTLs on non-gene chromosomes) as defined by the GENCODE [51] version 12 gene annotation. We calculated the proportion of the variance (narrow-sense heritability) explained by each component using the following mixed-effects model:

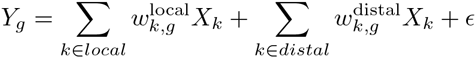

where *Y_*g*_* represents the expression of gene *g*, *X*_*k*_ is the allelic dosage for SNP *k*, local refers to the set of SNPs located within 1Mb of the gene's transcription start and end, distal refers to SNPs in other chromosomes, and e is the error term representing environmental and other unknown factors. We assume that the local and distal components are independent of each other as well as independent of the error term. We assume random effects for 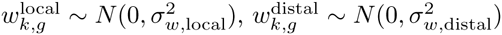, and 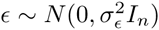, where *I*_*n*_ is the identity matrix. We calculated the total variability explained by local and distal components using restricted maximum likelihood (REML) as implemented in the GCTA software [28].

In an effort to determine if distal heritability estimates could be improved, we also tested a mixed-effect model restricting the distal component to known eQTLs on non-gene chromosomes discovered in the Framingham cohort at FDR < 0.05. When we found the distal estimate could not be improved, we focused on estimating local heritability without the distal component in the equation above.

For the purpose of estimating the mean heritability (see Table 1, S1 Fig and S1 Table), we allowed the heritability estimates to take negative values (unconstrained model). Despite the lack of obvious biological interpretation of a negative heritability, it is an accepted procedure used in order to avoid bias in the estimated mean [10, 29]. Genes with FDR < 0.1 (derived from the two-sided GCTA P-value) were considered to have significant heritability.

For comparing to BSLMM PVE, we restricted the GCTA heritability estimates to be within the [0,1] interval (constrained model, see Figures 2, 4 and 6).

### Quantifying sparsity with Bayesian Sparse Linear Mixed Modeling (BSLMM)

We used BSLMM [14] to model the effect of local genetic variation (common SNPs within 1 Mb of gene) on the genetic architecture of gene expression. BSLMM uses a linear model with a polygenic component (small effects) and a sparse component (large effects) enforced by sparsity inducing priors on the regression coefficients [14]. BSLMM assumes the genotypic effects come from a mixture of two normal distributions and thus is flexible to both polygenic and sparse genetic architectures [14]. We used the software GEMMA [54] to implement BSLMM for each gene with 100K sampling steps per gene. BSLMM estimates the PVE (the proportion of variance in phenotype explained by the additive genetic model, analogous to the heritability estimated in GCTA) and PGE (the proportion of genetic variance explained by the sparse effects terms where 0 means that genetic effect is purely polygenic and 1 means that the effect is purely sparse). From the second half of the sampling iterations for each gene, we report the median and the 95% credible sets of the PVE, PGE, and the |*γ*| parameter (the number of SNPs with non-zero coefficients).

### Determining polygenicity versus sparsity using the elastic net

We used the *glmnet* R package to fit an elastic net model where the tuning parameter is chosen via 10-fold cross-validation to maximize prediction performance measured by Pearson's R^2^ [52, 53].

The elastic net penalty is controlled by mixing parameter *α*, which spans LASSO (*α* = 1, the default) [19] at one extreme and ridge regression (*α* = 0) [20] at the other. The ridge penalty shrinks the coefficients of correlated SNPs towards each other, while the LASSO tends to pick one of the correlated SNPs and discard the others. Thus, an optimal prediction R^2^ for *α* = 0 means the gene expression trait is highly polygenic, while an optimal prediction R^2^ for *α* = 1 means the trait is highly sparse.

In the DGN cohort, we tested 21 values of the mixing parameter (*α* = 0, 0.05, 0.1, …, 0.90, 0.95, 1) for optimal prediction of gene expression of the 341 genes on chromosome 22. In order to compare prediction R^2^ values across *α* values, we used common folds and seeds for each run. For the rest of the autosomes in DGN and for whole tissue, cross-tissue, and tissue-specific expression in the GTEx cohort, we tested *α* = 0.05, 0.5, 1.

### Orthogonal Tissue Decomposition (OTD)

We use a mixed effects model to decompose the expression level of a gene into a subject-specific component and a subject-by-tissue-specific component. We fit the model one gene at a time and to simplify notation we assume the gene index, *g*, is implicit and drop it from the equations below. The expression *Y* of a gene *g* for individual *i* in tissue *t* is modeled as

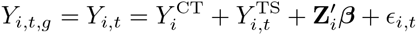

where 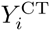 is the random subject level intercept, 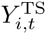 is the random subject by tissue intercept, Z_*i*_ represents covariates (for overall intercept, tissue intercept, gender, and PEER factors), and _*t*_ is the error term. We assume 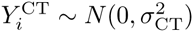, 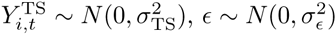, and that all three terms are independent of each other. All variances are estimated by restricted maximum likelihood (REML). *Y*_*i*,*t*,*g*_ = *Y*_*i*,*t*_ is a scalar. Z_*i*_ is a vector of length *p*, which represents the number of covariates. *β* is a vector of length *p* and represents the effects of covariates on the expression level of the gene. The prime in 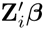 represents the inner product between the two vectors. All variances are estimated by restricted maximum likelihood (REML). These variances will be different for each gene. The tissue-specific component's variance is common across tissues. Differences between tissues will be reflected in the posterior mean of each tissue/individual random effects.

For the cross-tissue component 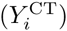 to be identifiable, multiple replicates of expression are needed for each subject. In the same vein, for the tissue-specific component 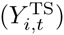 to be identifiable, multiple replicates of expression are needed for a given tissue/subject pair. GTEx [27] data consisted of expression measurement in multiple tissues for each subject, thus multiple replicates per subject were available. However, there were very few replicated measurement for a given tissue/subject pair. Thus, we fit the reduced model and use the estimates of the residual as the tissue-specific component.

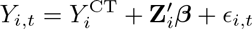

We consider the expression level of a gene at a given tissue for individual *i* to be composed of a cross-tissue component which we represent as 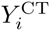 and a tissue-specific component, which given the lack of replicates, we estimate as the difference between the expression level and the cross-tissue components (after adjusting for covariates). The assumptions are the same as the full model, i.e., 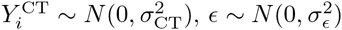, and that both terms are independent of each other.

The mixed effects model parameters were estimated using the `lme4 package` [55] in R. Batch effects and unmeasured confounders were accounted for using 15 `PEER` factors computed with the `PEER` [34] package in R. We also included gender as a covariate. Posterior modes of the subject level random intercepts were used as estimates of the cross-tissue components whereas the residuals of the models were used as tissue-specific components.

The model included whole tissue gene expression levels in 8555 GTEx tissue samples from 544 unique subjects. A total of 17,647 protein-coding genes (defined by GENCODE [51] version 18) with a mean gene expression level across tissues greater than 0.1 RPKM (reads per kilobase of transcript per million reads mapped) and RPKM > 0 in at least 3 individuals were included in the model.

### OTD simulation

To test the robustness of our method, we generated simulated expression levels so that we could compare our estimates with the ground truth. To preserve correlation structure between genes and tissues, we used the cross-tissue and tissue-specific estimates of OTD based on the observed data as a basis for the simulations. Thus, we simulated expression phenotypes as the sum of the observed OTD cross-tissue and tissue-specific components plus an error term derived by randomly sampling a normal distribution with a standard deviation equal to 1, 2, or 10 times the variance of each gene's component sum. We performed OTD on the simulated phenotypes and compared the OTD-estimated cross-tissue and tissue-specific expression levels for each gene to the true levels using Pearson correlation. As expected, higher levels of noise reduced the correlation between true and simulated values, but the correlation remained significant for all simulated phenotypes.

### Comparison of OTD trait heritability with multi-tissue eQTL results

To verify that the newly derived cross-tissue and tissue-specific traits were capturing the expected properties, we used the results of the multi-tissue eQTL analysis developed by Flutre et al. [33] and performed on nine tissues from the pilot phase of the GTEx project [27]. In particular, we downloaded the posterior probabilities of a gene being actively regulated in a tissue (PPA) from the GTEx portal at http://www.gtexportal.org/static/datasets/gtex_analysis_pilot_v3/multi_tissue_eqtls/Multi_tissue_eQTL_GTEx_Pilot_Phase_datasets.tar. PPA can be interpreted as the probability a gene is regulated by an eQTL in tissue *t* given the data. In the GTEx pilot, the most significant eQTL per gene was used to compute PPA [27]. PPA is computed from a joint analysis of all tissues and takes account of sharing of eQTLs among tissues [33]. For example, consider a SNP showing modest association with expression in tissue *t*. If this SNP also shows strong association in the other tissues, then it will be assigned a higher probability of being an active eQTL in tissue *t* than if it showed no association in the other tissues [33].

We reasoned that genes with large cross-tissue component (i.e. high cross-tissue h^2^) would have more uniform PPA across tissues. Thus we defined for each gene a measure of uniformity, *U*_*g*_, across tissues based on the nine-dimensional vector of PPAs using the entropy formula. More specifically, we divided each vector of PPA by their sum across tissues and computed the measure of uniformity as follows:

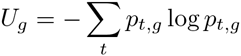

where *p*_*t*,*g*_ is the normalized PPA for gene *g* and tissue *t*.

## Acknowledgments

We thank Nicholas Knoblauch and Jason Torres for initial pipeline development and planning. We thank Nicholas Miller for assistance building the results database.

## GTEx data

The Genotype-Tissue Expression (GTEx) Project was supported by the Common Fund of the Office of the Director of the National Institutes of Health (commonfund.nih.gov/GTEx). Additional funds were provided by the NCI, NHGRI, NHLBI, NIDA, NIMH, and NINDS. Donors were enrolled at Biospecimen Source Sites funded by NCI Leidos Biomedical Research, Inc. subcontracts to the National Disease Research Interchange (10XS170), Roswell Park Cancer Institute (10XS171), and Science Care, Inc. (X10S172). The Laboratory, Data Analysis, and Coordinating Center (LDACC) was funded through a contract (HHSN268201000029C) to the The Broad Institute, Inc. Biorepository operations were funded through a Leidos Biomedical Research, Inc. subcontract to Van Andel Research Institute (10ST1035). Additional data repository and project management were provided by Leidos Biomedical Research, Inc.(HHSN261200800001E). The Brain Bank was supported supplements to University of Miami grant DA006227. Statistical Methods development grants were made to the University of Geneva (MH090941 & MH101814), the University of Chicago (MH090951,MH090937, MH101825, & MH101820), the University of North Carolina-Chapel Hill (MH090936), North Carolina State University (MH101819),Harvard University (MH090948), Stanford University (MH101782), Washington University (MH101810), and to the University of Pennsylvania (MH101822). The datasets used for the analyses described in this manuscript were obtained from dbGaP at **http://www.ncbi.nlm.nih.gov/gap** through dbGaP accession number phs000424.v3.p1.

## DGN data

NIMH Study 88 – Data was provided by Dr. Douglas F. Levinson. We gratefully acknowledge the resources were supported by National Institutes of Health/National Institute of Mental Health grants 5RC2MH089916 (PI: Douglas F. Levinson, M.D.; Coinvestigators: Myrna M. Weissman, Ph.D., James B. Potash, M.D., MPH, Daphne Koller, Ph.D., and Alexander E. Urban, Ph.D.) and 3R01MH090941 (Co-investigator: Daphne Koller, Ph.D.).

## Framingham data

The Framingham Heart Study is conducted and supported by the National Heart, Lung, and Blood Institute (NHLBI) in collaboration with Boston University (Contract No. N01-HC-25195 and HHSN268201500001I). This manuscript was not prepared in collaboration with investigators of the Framingham Heart Study and does not necessarily reflect the opinions or views of the Framingham Heart Study, Boston University, or NHLBI.

Funding for SHARe Affymetrix genotyping was provided by NHLBI Contract N02-HL-64278. SHARe Illumina genotyping was provided under an agreement between Illumina and Boston University. Funding for Affymetrix genotyping of the FHS Omni cohorts was provided by Intramural NHLBI funds from Andrew D. Johnson and Christopher J. O'Donnell.

Additional funding for SABRe was provided by Division of Intramural Research, NHLBI, and Center for Population Studies, NHLBI.

The following datasets were downloaded from dbGaP: phs000363.v12.p9 and phs000342.v13.p9.

## Computing resources

**OSDC** This work made use of the Open Science Data Cloud (OSDC) which is an Open Cloud Consortium (OCC)-sponsored project. This work was supported in part by grants from Gordon and Betty Moore Foundation and the National Science Foundation and major contributions from OCC members like the University of Chicago [57].

**Bionimbus** This work made use of the Bionimbus Protected Data Cloud (PDC), which is a collaboration between the Open Science Data Cloud (OSDC) and the IGSB (IGSB), the Center for Research Informatics (CRI), the Institute for Translational Medicine (ITM), and the University of Chicago Comprehensive Cancer Center (UCCCC). The Bionimbus PDC is part of the OSDC ecosystem and is funded as a pilot project by the NIH [58] (https://www.bionimbus-pdc.opensciencedatacloud.org/).

## Supporting Information

**S1 Fig.**
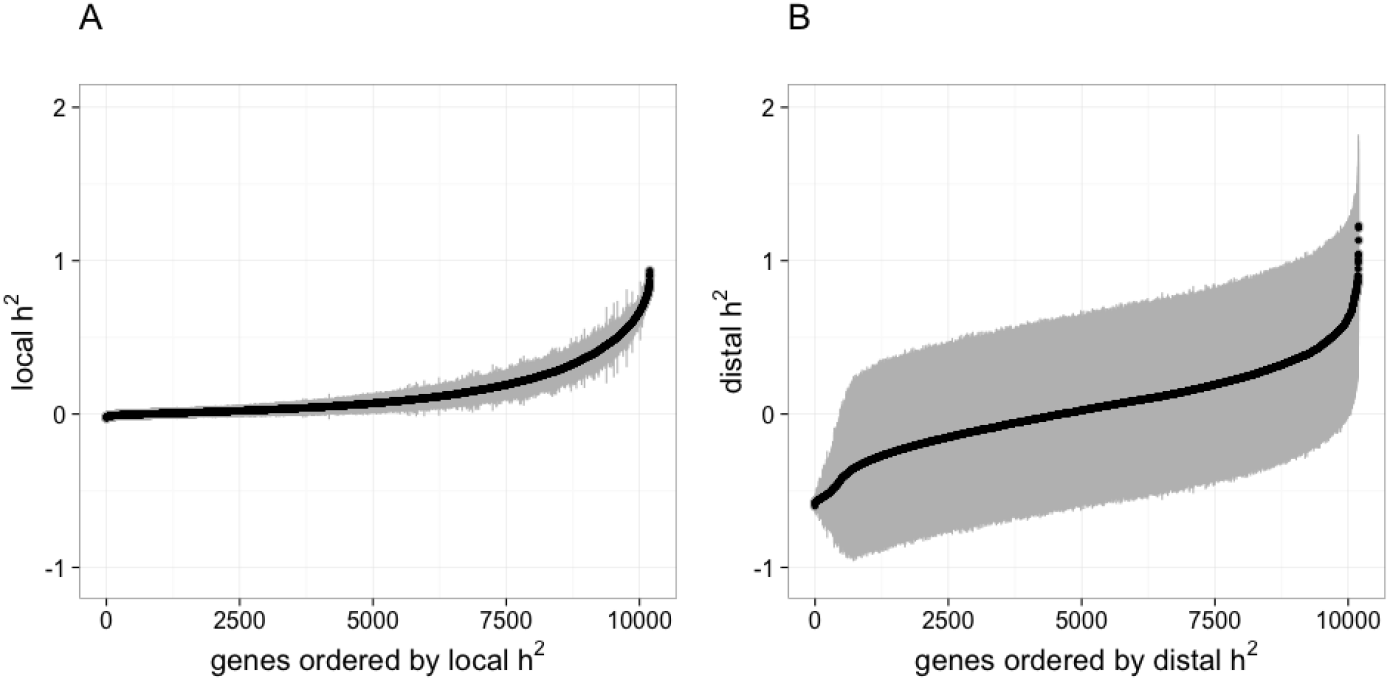
Local and distal heritability (h^2^) of gene expression levels in whole blood from the DGN RNA-seq dataset. In order to obtain an unbiased estimate of mean h^2^, we allow the values to be negative when fitting the restricted maximum likelihood (unconstrained REML). (A) Local (SNPs within 1Mb of each gene) and (B) distal (all SNPs on non-gene chromosomes) gene expression h^2^ estimates ordered by increasing h^2^. Gray segments show two times the standard errors of each h^2^ estimate. Notice the larger range and errors of the distal estimates.

**S2 Fig.**
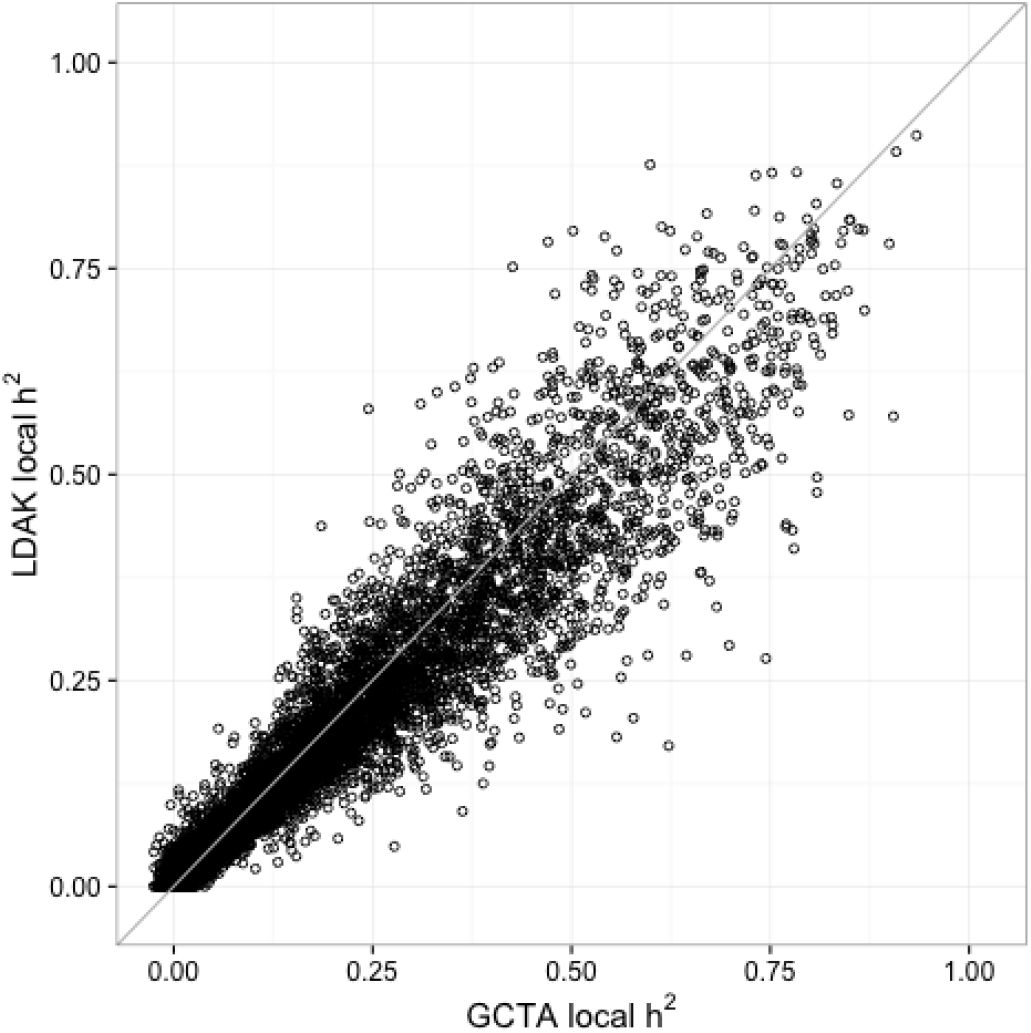
DGN local heritability estimation comparison between LDAK and GCTA. The Pearson correlation between GCTA and the LD-adjusted kinship local h^2^ estimates of gene expression is 0.96.

**S3 Fig.**
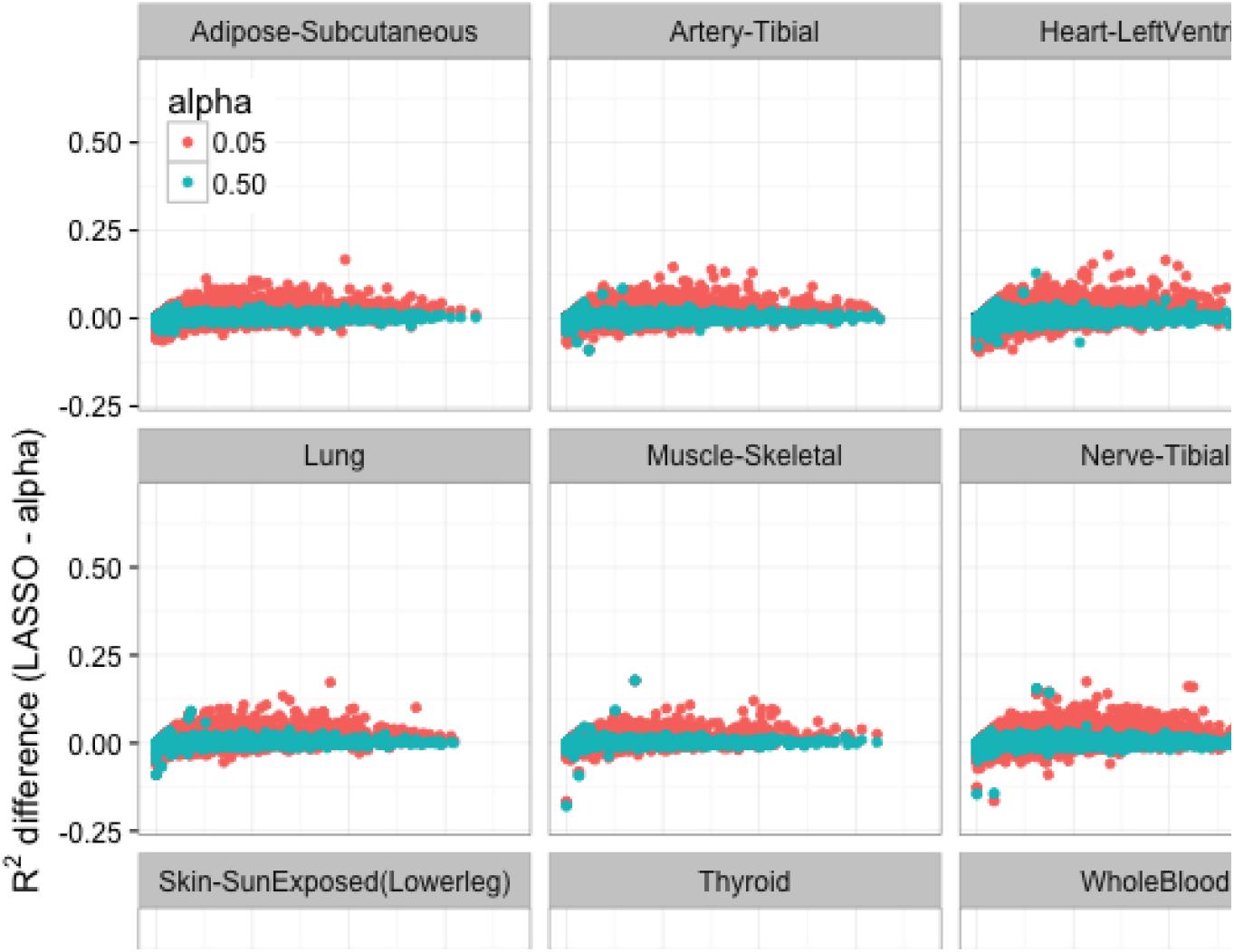
GTEx whole tissue cross-validated predictive performance across the elastic net. The difference between the cross validated R^2^ of the LASSO model and the elastic net model mixing parameters 0.05 and 0.5 for autosomal protein coding genes per tissue. Elastic net with *α* = 0.5 values hover around zero, meaning that it has similar predictive performance to LASSO. The R^2^ difference of the more polygenic model (elastic net with *α* = 0.05) is mostly above the 0 line, indicating that this model performs worse than the LASSO model across tissues.

**S4 Fig.**
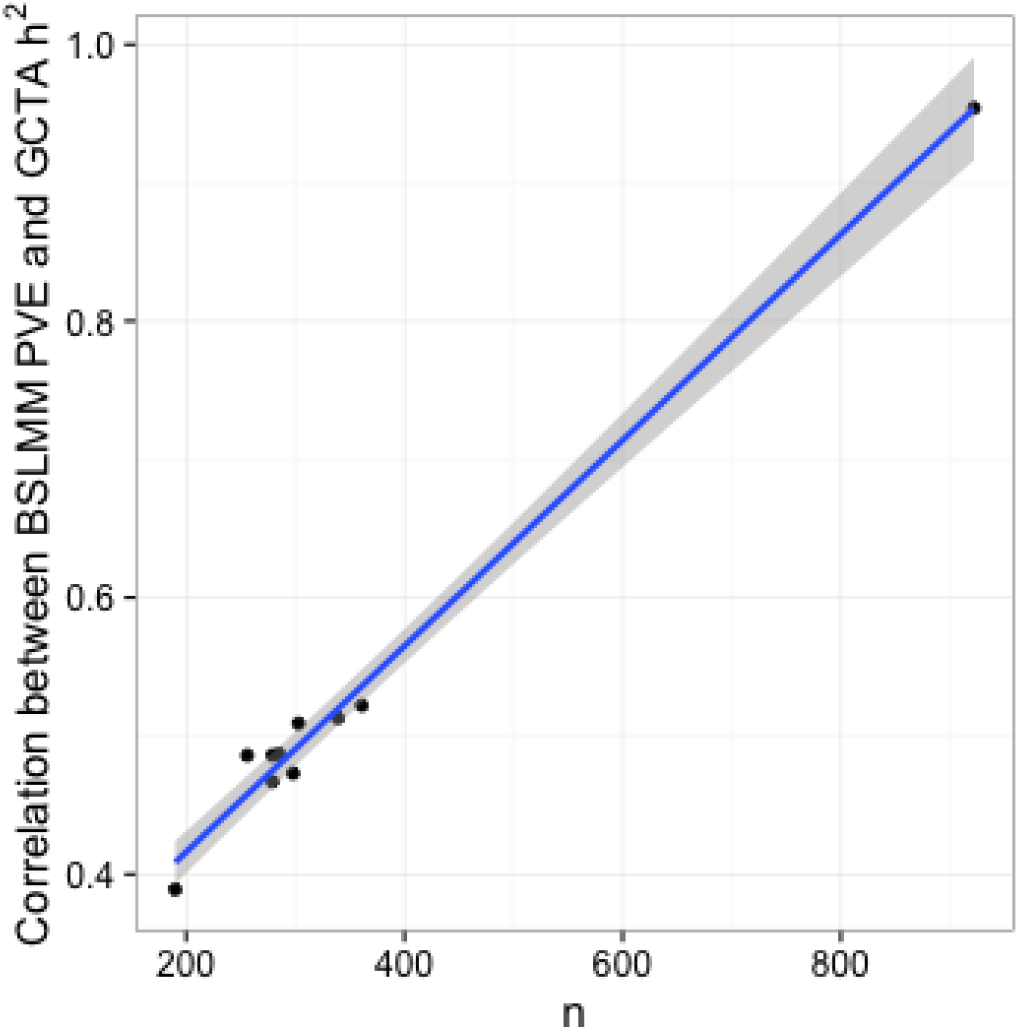
Correlation between heritability models compared to sample size. The Pearson correlation between estimates of heritability using BSLMM vs. GCTA is dependent on sample size (R = 0.99, *P* = 3*x*10^−9^).

**S5 Fig.**
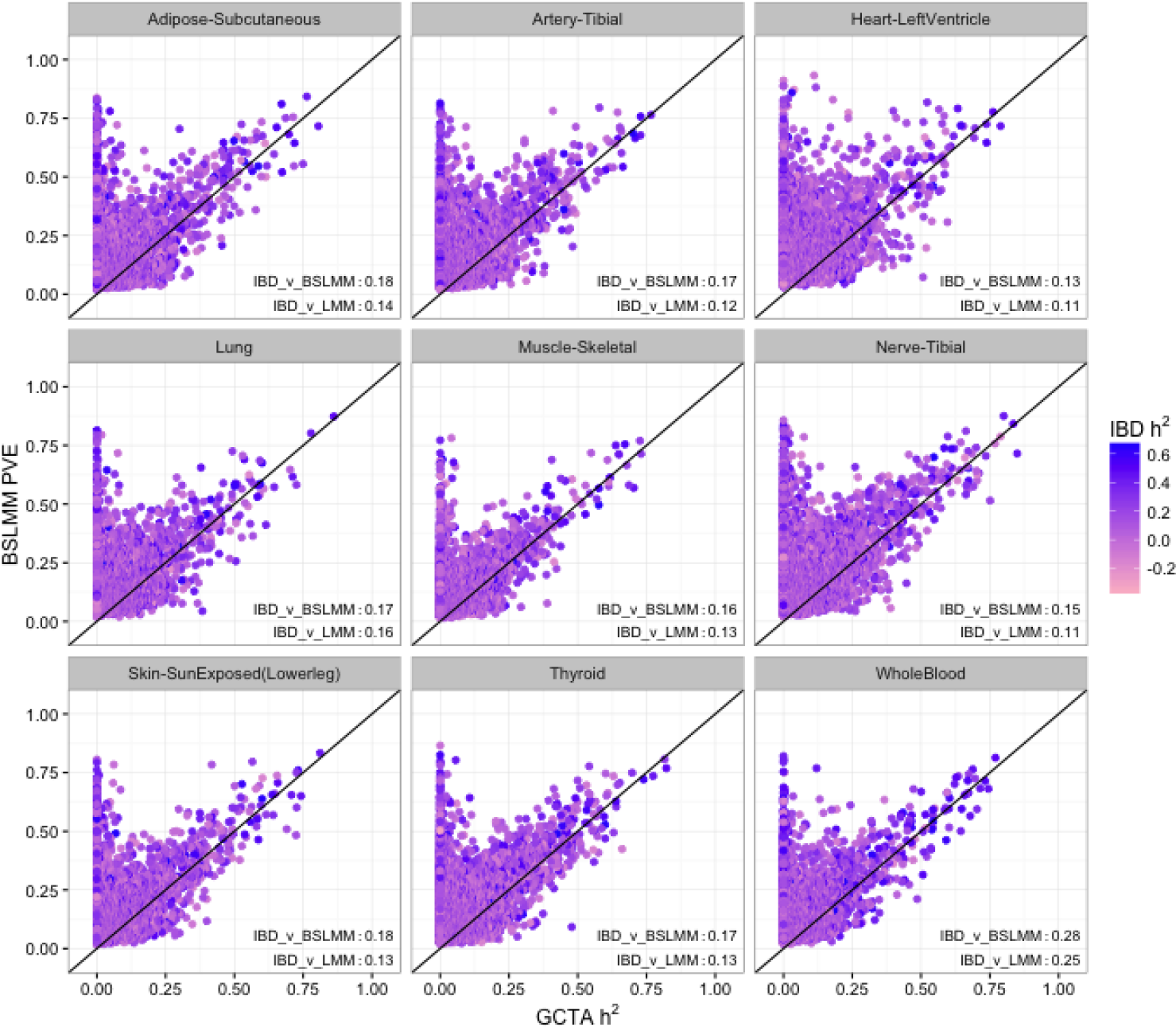
BSLMM and LMM estimates of heritability in GTEx compared to IBD estimates. This figure shows the comparison between estimates of heritability using BSLMM vs. LMM (GCTA) for GTEx data. Here, in both models the estimates are constrained to be between 0 and 1. For most genes BSLMM estimates are larger than LMM estimates reflecting the fact that BSLMM yields better estimates of heritability because of its ability to account for the sparse component. Each point is colored according to that gene's estimate of h^2^ by shared identity by descent (IBD) in Price et al. [29]. At the bottom left of each panel, we show the IBD correlation with BSLMM (IBD_v_BSLMM) and LMM (IBD_v_LMM). BSLMM is consistently more correlated with the IBD estimate. This provides further evidence that LMM is underestimating h^2^ for these genes.

**S6 Fig.**
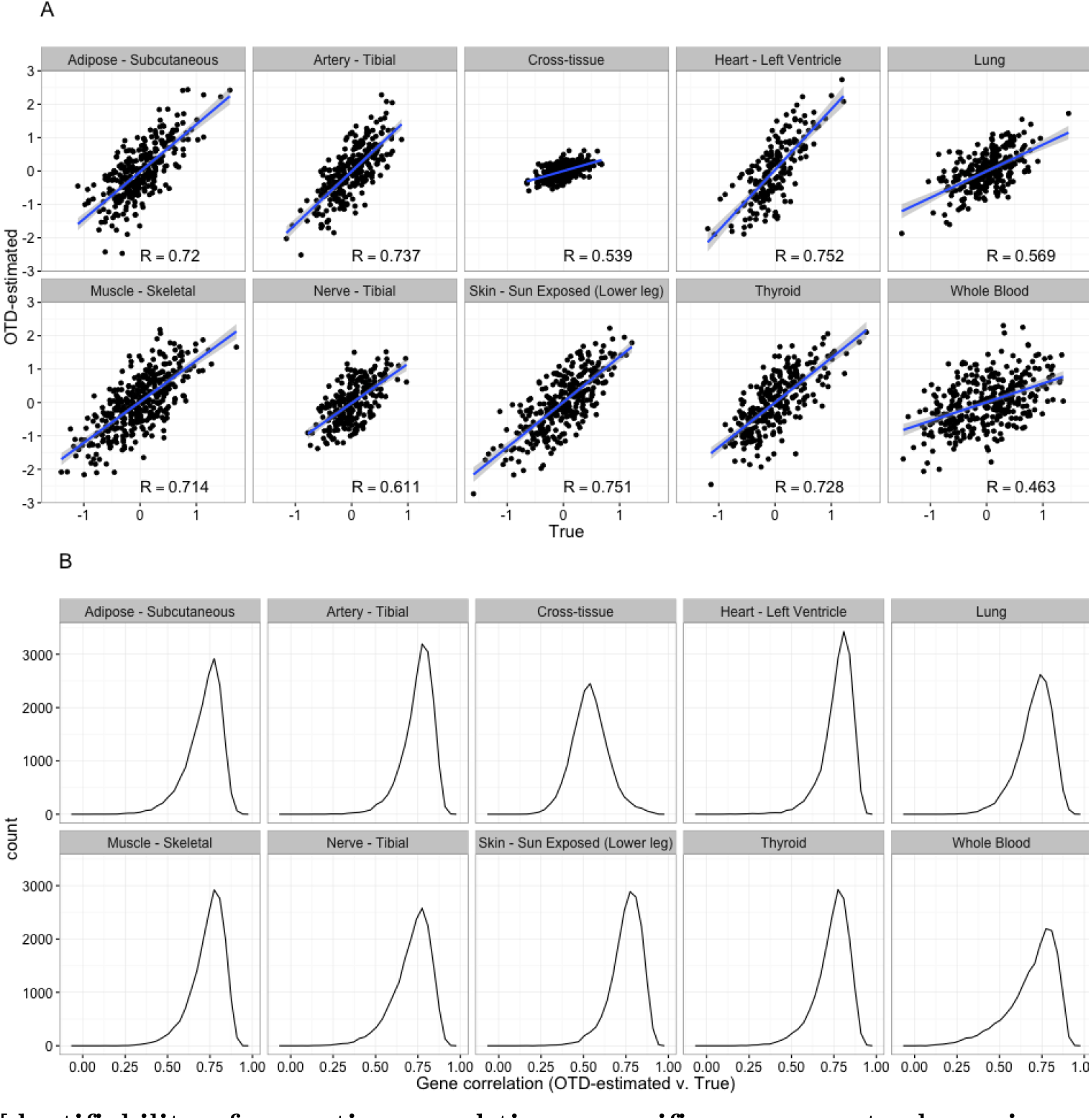
Identifiability of cross-tissue and tissue-specific components shown in simulated data. The simulated phenotype is the sum of the observed cross-tissue and tissue-specific components (estimated via OTD in the observed data) plus an error term derived by randomly sampling a normal distribution with variance equal to the variance of each gene. (A) Correlation between true and OTD-estimated cross-tissue and tissue-specific components for one gene, *SLTM*. (B) Histogram of the correlation between true and OTD-estimated components across all genes.

**S7 Fig.**
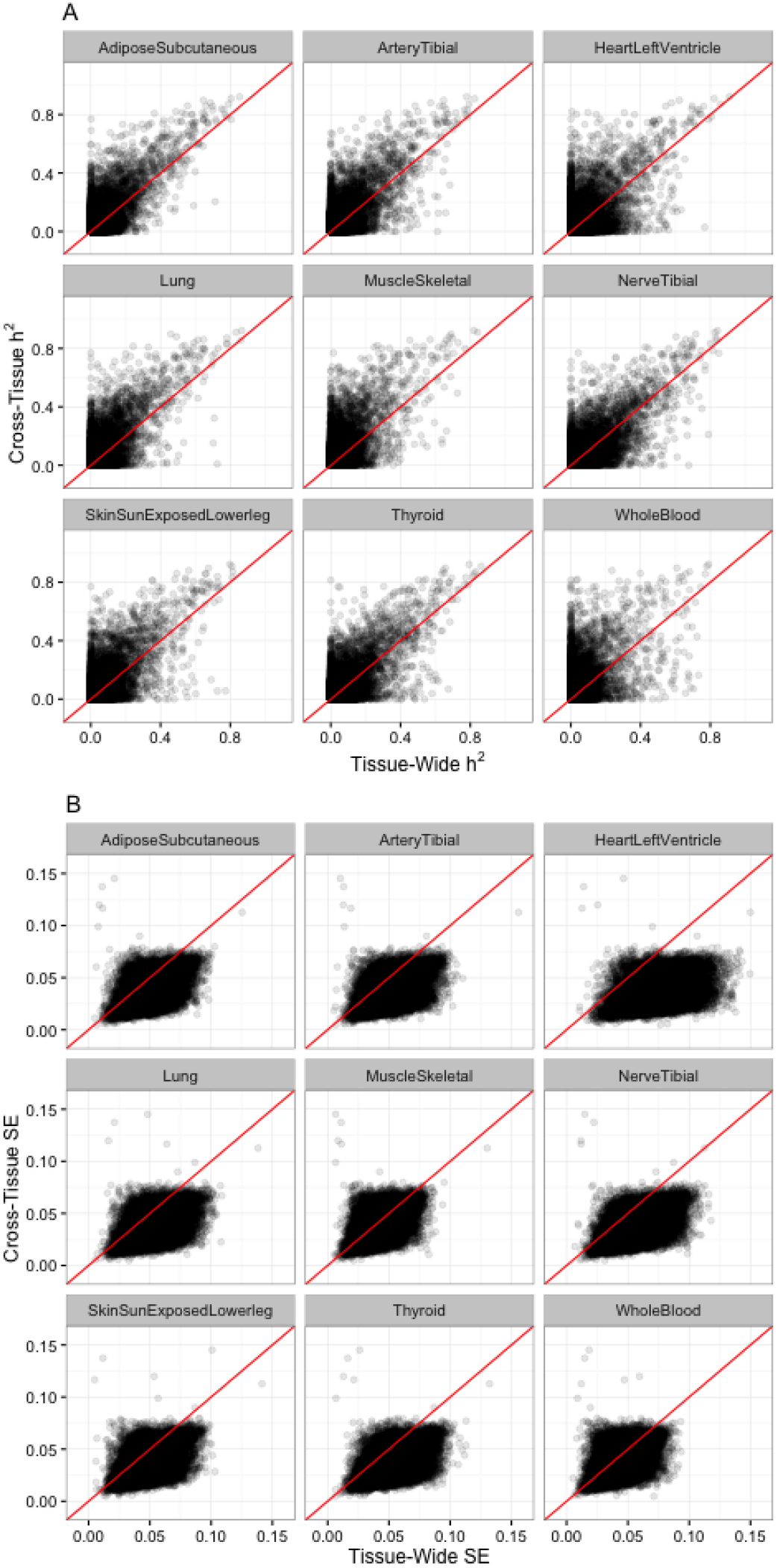
Cross-tissue and whole tissue comparison of heritability (h^2^, A) and standard error (SE, B). Cross-tissue local h^2^ is estimated using the cross-tissue component (random effects) of the mixed effects model for gene expression and SNPs within 1 Mb of each gene. Whole tissue local h^2^ is estimated using the measured gene expression for each respective tissue and SNPs within 1 Mb of each gene. Estimates of h^2^ for cross-tissue expression traits are larger and their standard errors are smaller than the corresponding estimates for each whole tissue expression trait.

**S8 Fig.**
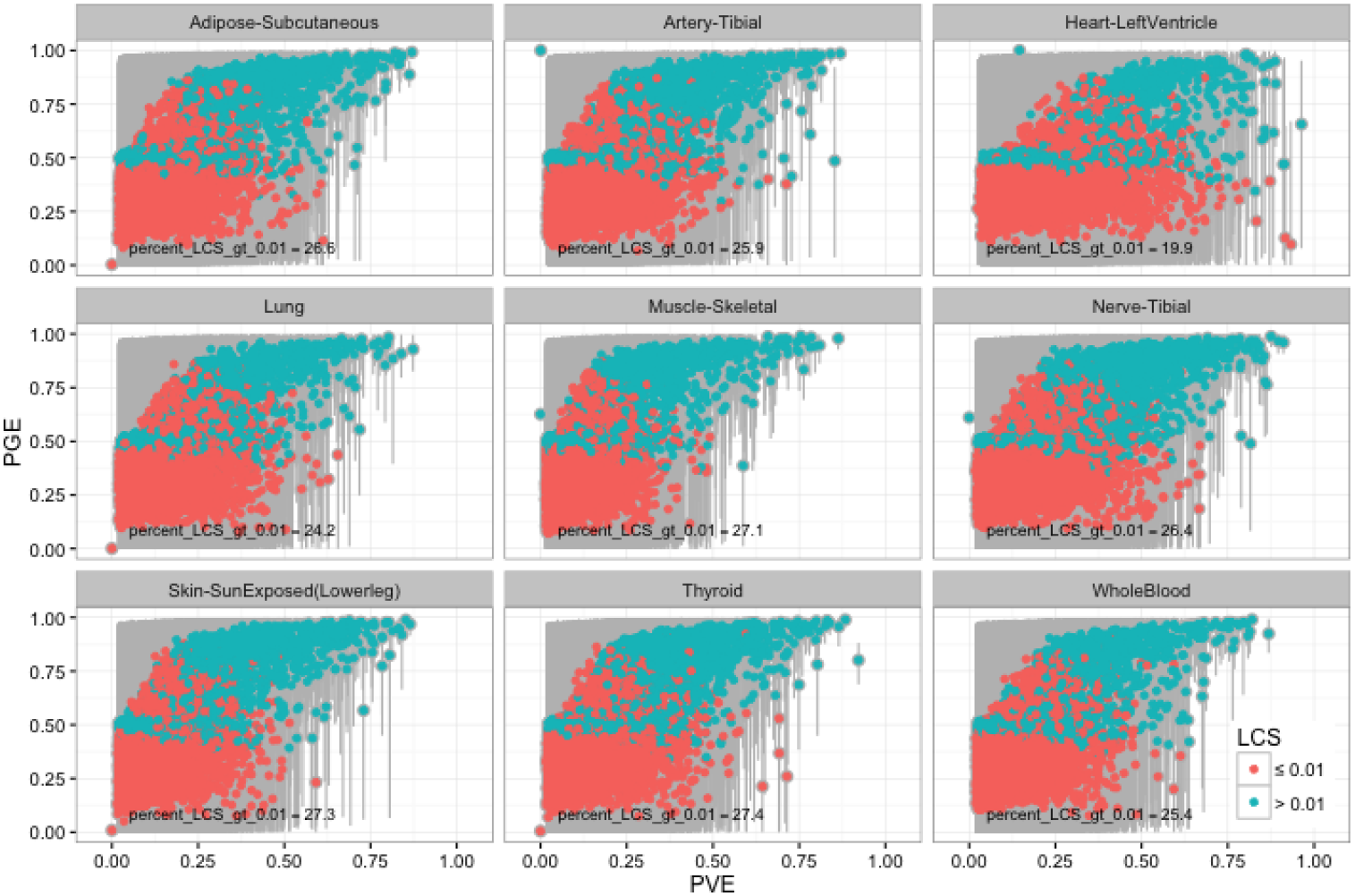
GTEx whole tissue expression Bayesian Sparse Linear Mixed Model. Comparison of median PGE (proportion of PVE explained by sparse effects) to median PVE (total proportion of variance explained, the BSLMM equivalent of h^2^) for expression of each gene. The 95% credible set of each PGE estimate is in gray and genes with a lower credible set (LCS) greater than 0.01 are in blue. For highly heritable genes the sparse component is close to 1, thus for high heritability genes the local architecture is sparse across tissues. For lower heritability genes, there is not enough evidence to determine sparsity or polygenicity.

**S9 Fig.**
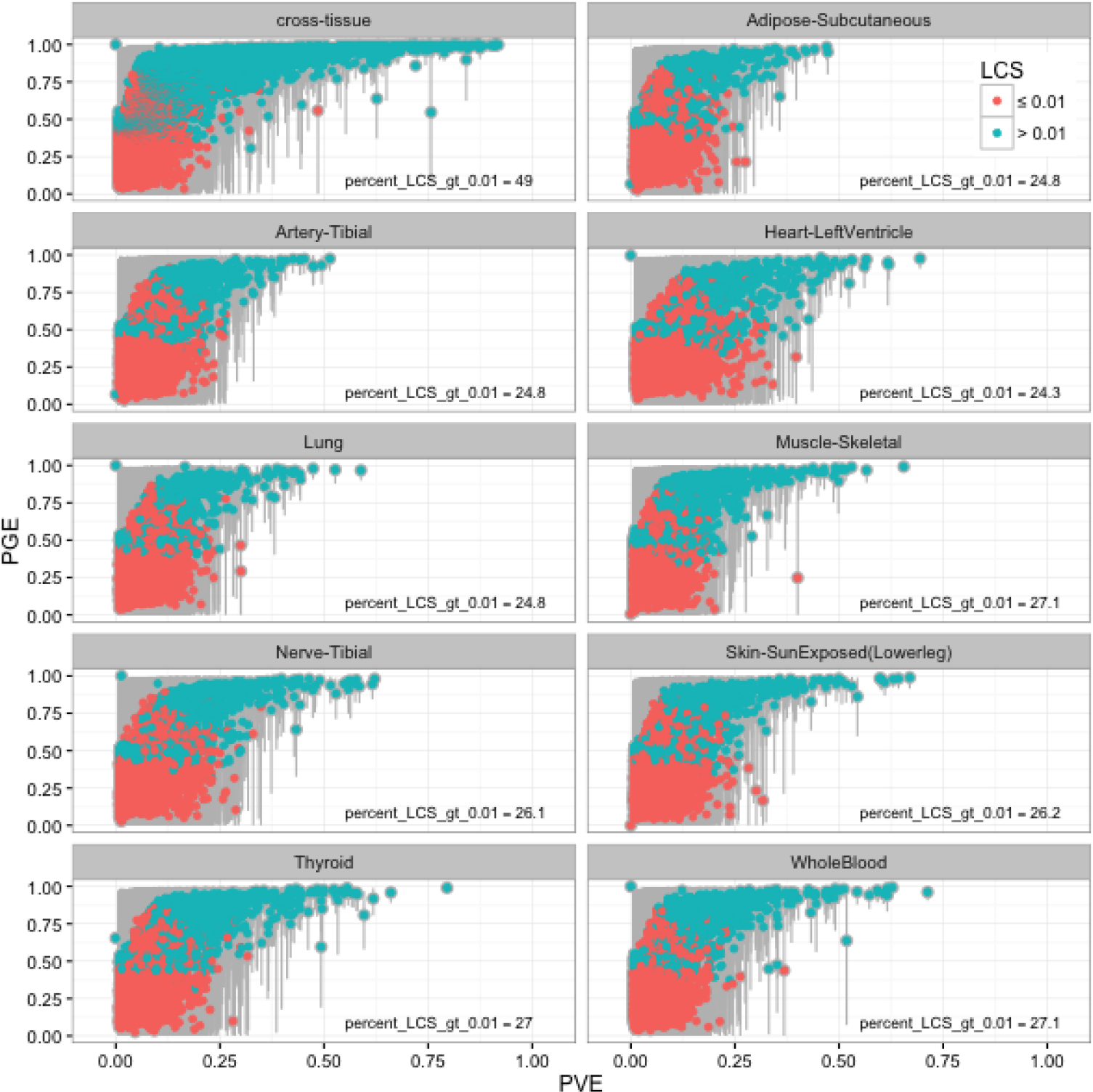
GTEx orthogonal tissue decomposition cross-tissue and tissue-specific expression Bayesian Sparse Linear Mixed Model. Comparison of median PGE (proportion of PVE explained by sparse effects) to median PVE (total proportion of variance explained, the BSLMM equivalent of h^2^) for expression of each gene. The 95% credible set of each PGE estimate is in gray and genes with a lower credible set (LCS) greater than 0.01 are in blue. For highly heritable genes the sparse component is close to 1, thus for high heritability genes the local architecture is sparse across tissues. About twice as many cross-tissue expression traits have significant PGE (LCS > 0.01) compared to the tissue-specific expression traits.

**S10 Fig.**
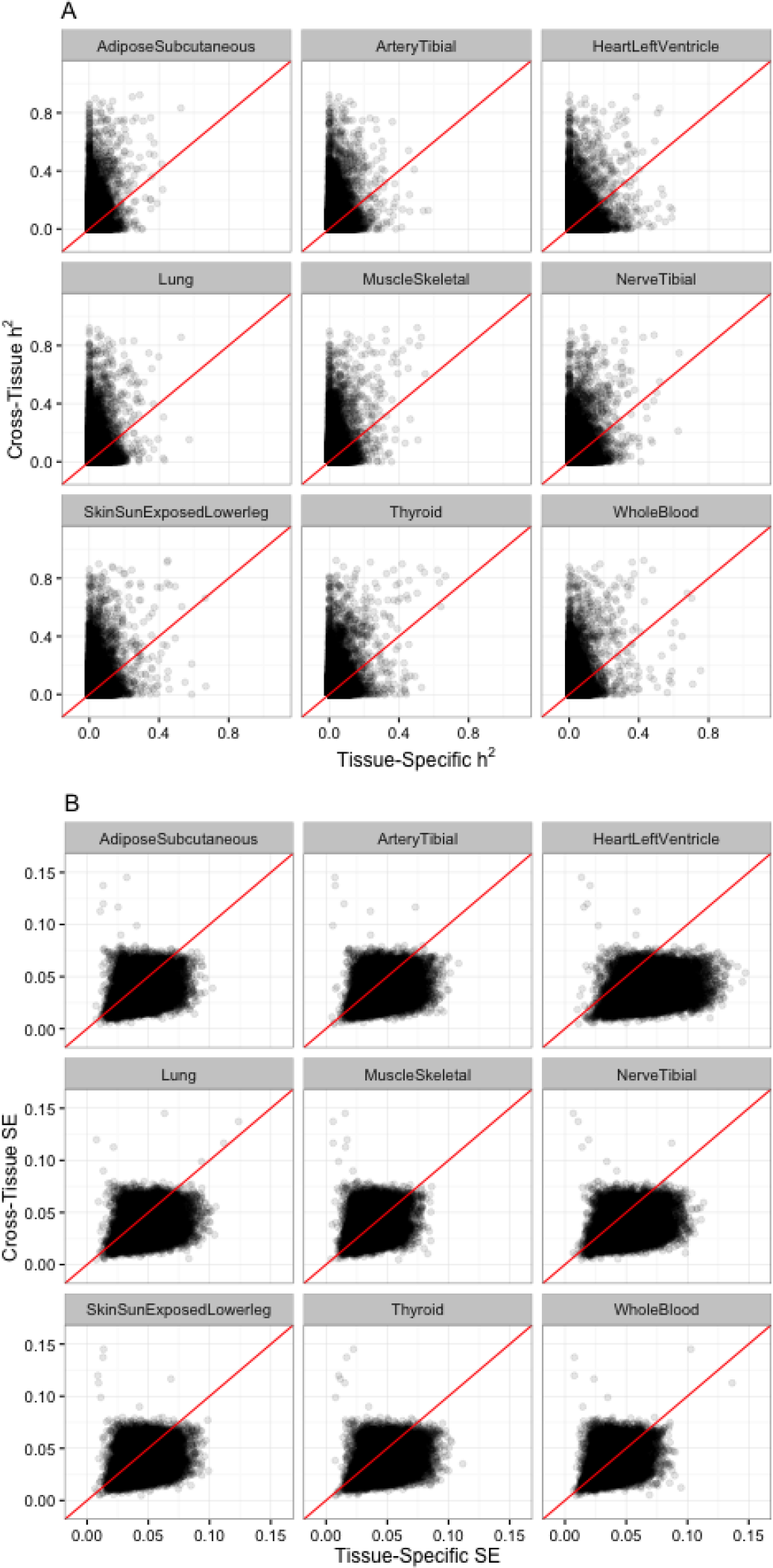
Cross-tissue and tissue-specific comparison of heritability (h^2^, A) and standard error (SE, B) estimation. Cross-tissue local h^2^ is estimated using the cross-tissue component (random effects) of the mixed effects model for gene expression and SNPs within 1 Mb of each gene. Tissue-specifc local h^2^ is estimated using the tissue-specific component (residuals) of the mixed effects model for gene expression for each respective tissue and SNPs within 1 Mb of each gene. Estimates of h^2^ for cross-tissue expression traits are larger and their standard errors are smaller than the corresponding estimates for each tissue-specific expression trait.

**S11 Fig.**
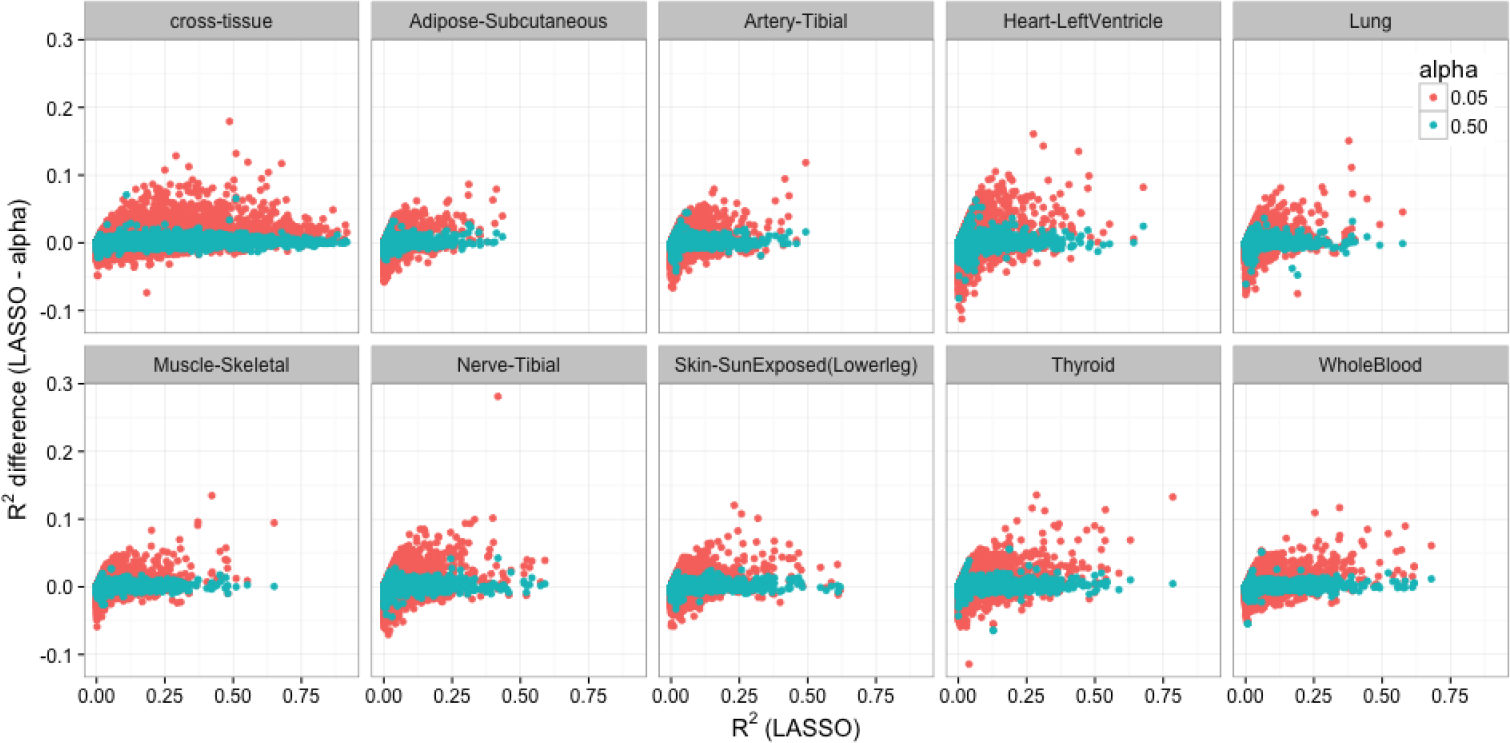
GTEx orthogonal tissue decomposition cross-tissue and tissue-specific expression cross-validated predictive performance across the elastic net. The difference between the cross validated R^2^ of the LASSO model and the elastic net model mixing parameters 0.05 and 0.5 for autosomal protein coding genes per cross-tissue and tissue-specific gene expression traits. Elastic net with *α* = 0.5 values hover around zero, meaning that it has similar predictive performance to LASSO. The R^2^ difference of the more polygenic model (elastic net with α = 0.05) is mostly above the 0 line, indicating that this model performs worse than the LASSO model across decomposed tissues.

**S12 Fig.**
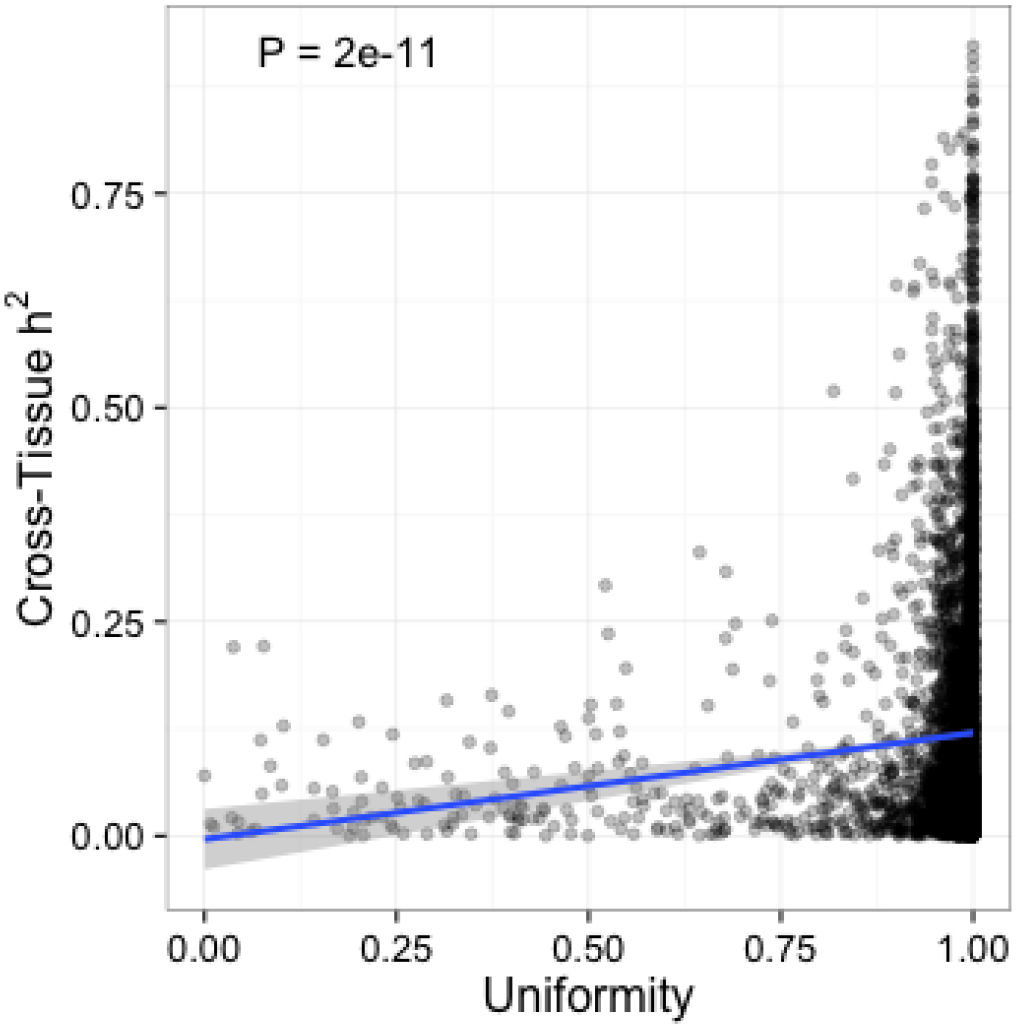
Continuous distribution of heritability of the cross-tissue component vs. a measure of uniformity of genetic regulation across tissues. The measure of uniformity was computed using the posterior probability of a gene being actively regulated in a tissue, PPA, from the Flutre et al. [33] multi-tissue eQTL analysis. Genes with PPA concentrated in one tissue were assigned small values of the uniformity measure whereas genes with PPA uniformly distributed across tissues were assigned high value of uniformity measure. See Methods for the entropy-based definition of uniformity. The blue line is the best fit linear regression and shows a significant positive correlation between h^2^ and uniformity (*β* = 0.12, *P* = 2e − 11).

**S1 Table.** Estimates of cross-tissue and tissue-specific local h^2^. Expression levels derived by Orthogonal Tissue Decomposition and h^2^ estimated using the `--reml-no-constrain` method.

| tissue | n | mean $h^2$ | % FDR<0.1 | num FDR<0.1 | num expressed |
| --- | --- | --- | --- | --- | --- |
| Cross-tissue | 450 | 0.062 | 20.2 | 2995 | 14861 |
| Adipose - Subcutaneous | 298 | 0.017 | 9.5 | 1408 | 14861 |
| Adrenal Gland | 126 | 0.037 | 8.9 | 1329 | 14861 |
| Artery - Aorta | 198 | 0.022 | 9.8 | 1449 | 14861 |
| Artery - Coronary | 119 | 0.048 | 8.1 | 1211 | 14861 |
| Artery - Tibial | 285 | 0.022 | 8.5 | 1262 | 14861 |
| Brain - Anterior cingulate cortex (BA24) | 72 | 0.037 | 10.9 | 1627 | 14861 |
| Brain - Caudate (basal ganglia) | 100 | 0.042 | 9.3 | 1388 | 14861 |
| Brain - Cerebellar Hemisphere | 89 | 0.046 | 10.9 | 1627 | 14861 |
| Brain - Cerebellum | 103 | 0.041 | 10.9 | 1618 | 14861 |
| Brain - Cortex | 96 | 0.053 | 10.0 | 1480 | 14861 |
| Brain - Frontal Cortex (BA9) | 92 | 0.037 | 10.6 | 1569 | 14861 |
| Brain - Hippocampus | 81 | 0.033 | 10.9 | 1618 | 14861 |
| Brain - Hypothalamus | 81 | 0.020 | 12.4 | 1839 | 14861 |
| Brain - Nucleus accumbens (basal ganglia) | 93 | 0.040 | 10.7 | 1585 | 14861 |
| Brain - Putamen (basal ganglia) | 82 | 0.033 | 10.6 | 1581 | 14861 |
| Breast - Mammary Tissue | 183 | 0.020 | 9.2 | 1367 | 14861 |
| Cells - EBV-transformed lymphocytes | 115 | 0.045 | 8.9 | 1323 | 14861 |
| Cells - Transformed fibroblasts | 272 | 0.019 | 9.0 | 1334 | 14861 |
| Colon - Sigmoid | 124 | 0.024 | 10.6 | 1578 | 14861 |
| Colon - Transverse | 170 | 0.022 | 10.6 | 1573 | 14861 |
| Esophagus - Gastroesophageal Junction | 127 | 0.029 | 9.1 | 1358 | 14861 |
| Esophagus - Mucosa | 242 | 0.026 | 8.2 | 1220 | 14861 |
| Esophagus - Muscularis | 219 | 0.024 | 9.0 | 1344 | 14861 |
| Heart - Atrial Appendage | 159 | 0.031 | 9.7 | 1437 | 14861 |
| Heart - Left Ventricle | 190 | 0.025 | 8.9 | 1325 | 14861 |
| Liver | 98 | 0.036 | 10.2 | 1519 | 14861 |
| Lung | 279 | 0.018 | 8.8 | 1308 | 14861 |
| Muscle - Skeletal | 361 | 0.020 | 9.0 | 1342 | 14861 |
| Nerve - Tibial | 256 | 0.026 | 8.3 | 1235 | 14861 |
| Ovary | 85 | 0.043 | 10.2 | 1523 | 14861 |
| Pancreas | 150 | 0.036 | 9.4 | 1396 | 14861 |
| Pituitary | 87 | 0.044 | 10.0 | 1481 | 14861 |
| Skin - Not Sun Exposed (Suprapubic) | 196 | 0.041 | 6.3 | 938 | 14861 |
| Skin - Sun Exposed (Lower leg) | 303 | 0.027 | 6.9 | 1020 | 14861 |
| Small Intestine - Terminal Ileum | 77 | 0.046 | 11.7 | 1733 | 14861 |
| Spleen | 89 | 0.062 | 9.7 | 1448 | 14861 |
| Stomach | 171 | 0.020 | 10.3 | 1532 | 14861 |
| Testis | 157 | 0.042 | 10.5 | 1561 | 14861 |
| Thyroid | 279 | 0.024 | 8.7 | 1293 | 14861 |
| Whole Blood | 339 | 0.025 | 7.7 | 1150 | 14861 |

All tissues are from the GTEx Project. Mean heritability ( $h^2$ ) is calculated across genes for each tissue. The percentage (%) and number (num) of genes with significant $h^2$ estimates (FDR < 0.1) in each tissue are reported.

